# Linking structure to function in high performing electrosynthetic biofilm communities

**DOI:** 10.64898/2025.12.16.694578

**Authors:** Rozanne Stroek, Minke Gabriëls, Marijn Winkelhorst, Marcel van den Broek, Martin Pabst, Ludovic Jourdin, Jean-Marc Daran, Djordje Bajic

## Abstract

Biofilm-based microbial electrosynthesis (MES) is a promising technology that converts CO2 into industrially relevant organic compounds using renewable energy sources. The highest performing MES systems reported to date consist of biofilm-driven microbial communities. However, for a successful deployment of this technology, we still need to overcome key challenges, including the long colonization time of the biocathode and the difficulty of controlling the product profile. To address these challenges, it is crucial to better understand the key microbial components responsible for the desired metabolic products, and how they assemble and function as a community. In this study, we conducted an in-depth characterisation of three high-performing mixed MES communities using metagenomics, metaproteomics and advanced analysis of metagenome-derived metabolic networks. Our findings identified Eubacterium limosum, Sporomusa sphaeroides and Clostridium aromativorans as key contributors to the production of acetate, butyrate and caproate via the Wood-Ljungdahl and the reverse β-oxidation pathways. A higher production of butyrate and caproate was observed in reactors with higher abundance of C. aromativorans, an organism only recently discovered and never reported in gas-fermenting systems before. The recovered genome of C. aromativorans, reconstructed in a single circular fragment, provides a more comprehensive genomic representation than the current reference. In addition, we found genes related to lactate and ethanol production from acetyl-CoA in the metagenomes, including proteomic evidence for lactate production. This study provides key insights into the microbial players, metabolic processes and microbial community features driving product formation in biofilm-based MES, bringing this technology closer to industrial application.

## Introduction

Climate change is imposing an increasingly severe strain on our environment, creating an urgent need for societal and technological alternatives to current unsustainable practices. Microbial biotechnologies are emerging as a key tool in this transition [1]. Microbial systems can enhance soil carbon sequestration [2,3], enable sustainable agricultural management [4–6], reduce methane emissions [7], convert waste into biogas and biofuels [8,9], and generate bioplastics during wastewater treatment [10]. To rationally design and optimize such applications, it is essential to develop a predictive, system-level understanding of microbial systems that integrates the rapidly expanding wealth of multi-omics data. Achieving this vision promises to unlock the vast metabolic potential of microbes and lay the foundations for the rational engineering of microbial systems as powerful, programmable tools across diverse applications.

Modern industrial biotechnologies rely largely on workhorse organisms such as *Saccharomyces cerevisiae* or *Escherichia coli*, because of their genetic and physiological accessibility. However, the use of individual organisms presents some important limitations in their ability to process complex substrates [11], product profile diversity [12] and titers [13], and evolutionary stability [14,15]. Multi-species microbial communities are increasingly recognized as an alternative that could help us overcome these limitations, expanding and improving the performance of individual strains [16]. Microbial communities are well known for their suitability to process complex [17], variable [18] or recalcitrant substrates [19], for example in fields such as wastewater treatment. Increasingly, there is also an interest in synthetic multi-species communities, which could enable the diversification of the product profile and titers through division of labor [20] and offer increased resistance to contamination through more complete occupation of available metabolic niches [21]. The potential for synergistic interactions between species has been also hypothesized to enable the use of simpler, more cost-effective media [22,23]. More broadly, microbial communities may allow us to access the metabolic capabilities of microbes that cannot be cultured in isolation [24,25]. Despite these promises, the use of microbial communities in many industries is still in its infancy due to our limited understanding of how interactions between community members shape their composition, stability and function.

Historically, our understanding of microbial communities has been hampered by their limited experimental tractability. The presence of uncultured species [26], the complex web of ecological interactions across genetic, physiological, and evolutionary levels, and their often intricate spatial structures pose major challenges to mechanistic characterization. In recent decades, multi-omics technologies have begun to bridge this gap, providing unprecedented insight into the structure and function of multi-species microbial systems [27]. Yet, these analyses are typically fragmented: genomic, transcriptomic, proteomic, and metabolomic data are often generated in separate experiments, by different groups, and published in isolation [28]. In addition, omics studies most often lack an integration of data into predictive models, such as (meta)genome-scale metabolic models [29]. Overall, this fragmentation hinders the development of truly predictive data-driven frameworks for community function.

Here, we address this challenge by integrating state-of-the-art omics techniques, including genomics and proteomics and embedding them directly into predictive meta–genome-scale metabolic models. We focus on microbial electrosynthesis (MES) communities, a promising biotechnology that leverages the ability of microbes to reduce CO₂ into valuable chemicals using externally supplied electrical current. While significant advances in electrode and reactor engineering have rapidly improved MES productivity [30], the microbial consortia that drive these processes have remained largely uncharacterized.

A typical MES reactor consists of two chambers separated by a cation exchange membrane; one containing an anode, and the other a cathode. In the anodic chamber, water is split to oxygen and protons, releasing electrons to the anode. These electrons require an energy input to elevate their energy levels, such as power supplied from green electricity. While protons diffuse across the membrane into the cathodic chamber, oxygen does not, ensuring an anaerobic environment. In the cathodic chamber, microorganisms in suspension or within a biofilm-covered cathode utilize these electrons to grow and reduce CO_2_ to short- and medium-chain carboxylic acids (MCCA’s) **(Figure 1**). The feasibility of MES was first demonstrated by Nevin et al. (2010) who used an acetogen, *Sporomusa ovata*, to synthesize extracellular multicarbon compounds (acetate and 2-oxobutyrate) from CO_2_ [31]. Subsequently, the field shifted towards the use of mixed microbial communities that, in general, have an increased product performance [32] and that colonize the cathode [33–38]. These biofilm-based reactors provide several important advantages, including their long-term stability, the absence of biocatalyst washout and the possibility of continuous operation [39]. Additionally, the use of mixed cultures enables the diversification of the product profile through division of labour and the use of simpler, more cost-effective media due to potential synergistic interactions between species. Several studies have expanded the MES product profile to include short and medium-chain fatty acids, alcohols, methane, and hydrogen [36,40–43]. MCCA’s serve as platform chemicals for the production of e.g. plastics, pharmaceuticals, and cosmetics [44] while alcohols such as ethanol can be refined into biofuels or used as substrate for fermentations to produce protein-rich biomass [45,46]. Methane and hydrogen are considered less desirable products due to their lower market values [47].

**Figure 1.**
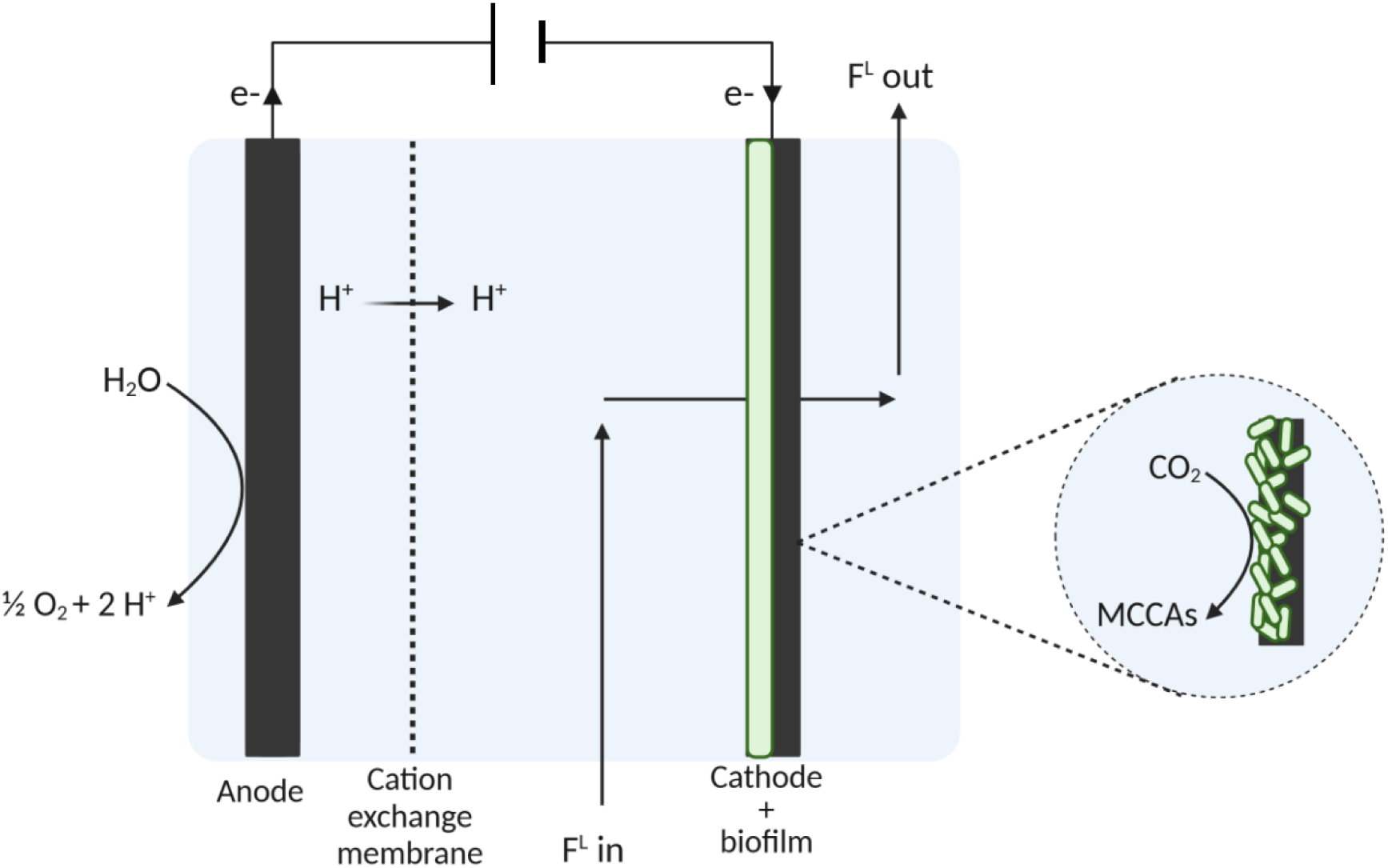
Schematic representation of a biofilm-based MES reactor. In the anodic chamber, water is oxidized to O2 and protons, releasing electrons to the anode. The electrons are flowing to the cathode side with power supplied by e.g. green energy. In the cathodic chamber, microbes on the biofilm-covered cathode convert CO_2_ into medium chain carboxylic acids, with the cathode serving as the electron donor.

Over the years, significant advancements in electrode and reactor engineering have been made to improve product titres in MES reactors, mostly focusing on cathode material development and reactor design [35,39,41,48–56]. These improvements have enabled MES systems to achieve volumetric productivities comparable to commercial syngas fermentation [39]. However, much less work has been done to characterize the composition of MES biofilm communities and the metabolic interactions dictating their assembly and function [30,57]. A better understanding of the biology of these communities could allow us to overcome important hurdles that currently prevent the industrial application of MES, such as the prolonged cathode colonisation time (100-225 day [39,58]), and the lack of control over product profile and specificity. Characterizing the communities at the core of MES bioreactors is essential for their manipulation and engineering. For example, it has been argued that establishing efficient microbe-electrode interactions and optimising mass transport across the electron-biofilm-environment interface are critical to achieving high productivity [59]. More broadly, understanding these communities might open the path towards the design of synthetic communities (SynComs) in MES, potentially enabling better control compared to mixed cultures [16].

Previous characterisations of biofilm-based MES communities have primarily relied on 16S rRNA gene sequencing, consistently identifying genera such as *Acetobacterium*, *Desulfovibrio*, *Arcobacter*, and *Sulfurospirillum* [35–37,43,55,60,61]. Whilst providing insights into the community composition, these studies have offered little information about the metabolic functions that drive growth, product formation and interactions between community members. Only a handful of studies have gone further. Vassilev et al. used a metagenomics approach to identify genes involved in carboxylic acid production, identifying *Clostridia* and *Rummeliibacillus* as dominant taxa in their MES reactors [43]. Marshall *et al.* reconstructed genome – scale models to identify key metabolic reactions underlying acetate production [62]. While these studies provide valuable insights, they are limited to either the metagenomic analysis of communities, thus lacking functional information, or solely acetate-producing communities.

Recently, Winkelhorst *et al.* reported MES reactors achieving high acetate, butyrate and caproate production rates [58]. Here, we characterize in depth the communities operating in these reactors by integrating high resolution metagenomics and metaproteomics with cutting edge metabolic network reconstruction and analysis algorithms. We identify key microbial players, reconstruct their genomes, and pinpoint the metabolic pathways that likely underlie distinct product profiles. Our analysis also identifies putative auxotrophies of key community members and potential cross-feeding interactions. Our results pave the way for the rational optimization of microbial electrosynthetic biofilm communities and delineate a broadly applicable path to the characterization of structure-function relationships in microbial communities.

## Results

In 2023, Winkelhorst *et al*. described the performance of four anaerobic electrochemical reactors, run as replicates, but producing distinctly different product profiles. The microbial communities of three of the four reactors were investigated in this study (reactor 1 (R1), reactor 2 (R2) and reactor 4 (R4)) [58]. During the last 20 days of operation, all three reactors produced acetate and butyrate, with reactor 4 exhibiting the highest acetate and butyrate concentration and rate. Caproate was additionally produced in reactor 1 and 4 **(Figure 2A&2B)**. Metagenomic and -proteomic samples were taken from each reactor’s biofilm at the last day of operation. This study describes an in-depth investigation into the microbial communities within these three reactors to gain a deeper understanding of the performance distinctions between them **(Supplemental information 1; Table S1).**

**Figure 2.**
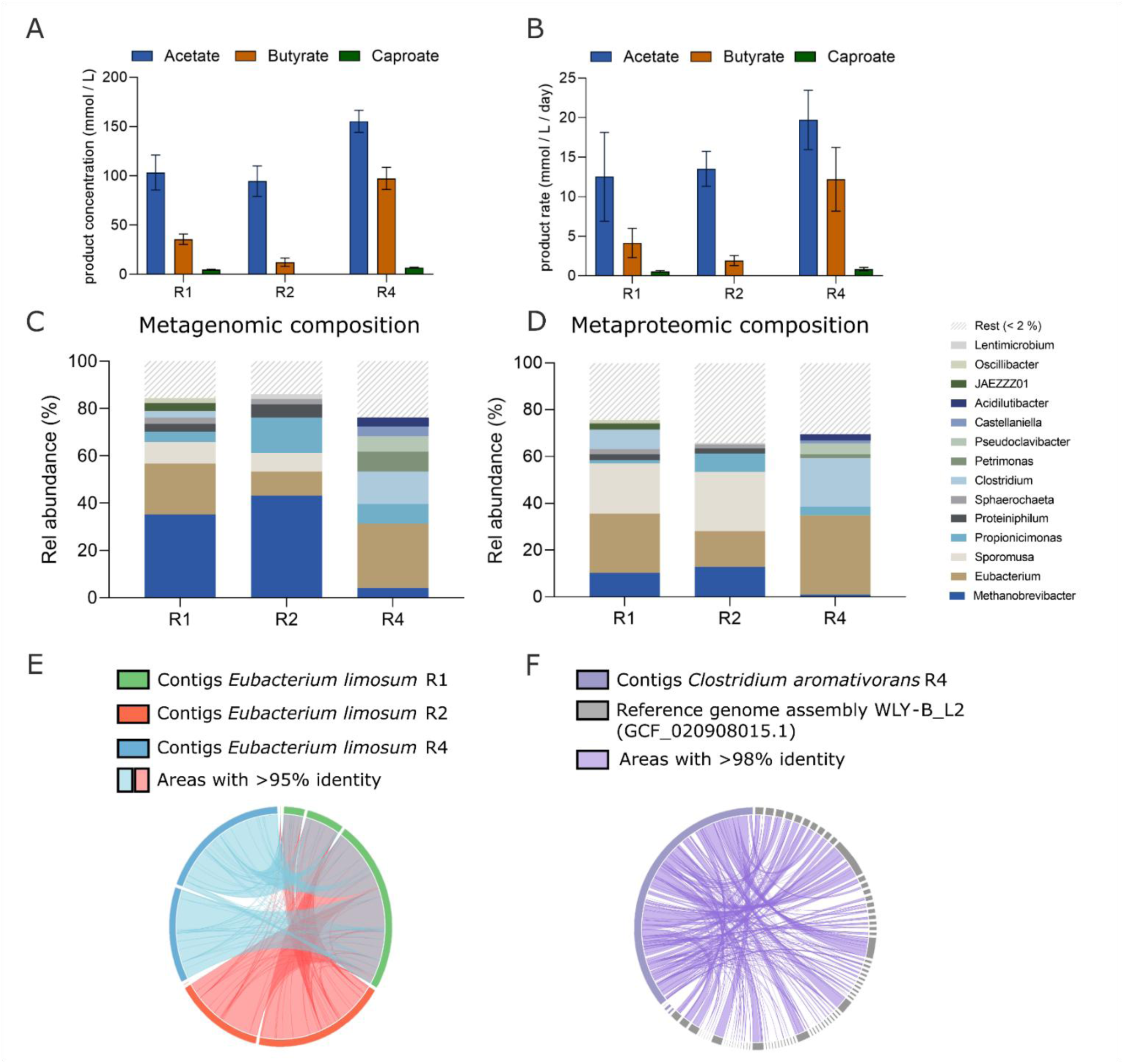
Mixed microbial community composition of three acetate, butyrate and caproate producing microbial electrosynthesis (MES) reactors. A & B: MES reactor performance during the last 20 days of operation of reactors 1, 2, and 4 (R1, R2 and R4) from the study done by Winkelhorst et al [58], including the acetate, butyrate and caproate concentration (A) and production rate (B). Acetate production is indicated in blue, butyrate production in orange, and caproate in green. C & D: Relative abundance of isolated metagenome-assembled genomes (MAGs) (genus level) within the microbial communities of three MES reactors as according to metagenomic sequencing data using long-read nanopore sequencing, polished with Illumina short-read sequencing (left), and metaproteomic data (right). “Rest” describes those sequences which did not meet the quality standards of CheckM2, were unbinned, or fell under a 2% relative abundance threshold. Metagenomic data was based on single samples. Metaproteomic data was combined of three samples taken at different locations of the biofilm. E. MAGs taxonomically identified to be Eubacterium limosum species in each of the reactors were aligned using BLASTn and areas of the genomes were colored based on >95% identity and e-value <1e-50 (light blue and light red). The MAG of reactor 1 is shown in green, reactor 2 in red and reactor 4 in blue. F. BLASTn alignment of as Clostridium aromativorans taxonomically identified MAG in R4 (purple) to the current reference genome of C. aromativorans (grey) from the study of Luo et al., 2023. The areas of the genome were colored based on >98% identity and e-value< 1e-50 (light purple) (see Supplemental information 2 and 3 for alignments of other MAGs).

### Conserved microbial community composition in Reactors 1 and 2, greater diversity within Reactor 4

To shed light on the composition of the microbial communities within the reactors, biofilm samples were collected from the cathodes after 194 days of operation and analysed using both whole genome sequencing (WGS) and 16S rRNA gene sequencing. Although 16S rRNA gene sequencing is a well-established method for analysing mixed communities [35–37,43,55,60,61], we observed notable disparities between the results obtained from the two methods (**Supplemental information 1; Figure S1**). Given the greater reliability of WGS for taxonomic profiling, community composition was based on WGS data. Fourteen genera were identified across the three reactors, with a consistent presence seen for *Methanobrevibacter*, *Eubacterium* and *Propionicimonas* in each reactor **(Figure 2C)**. The complexity of the community varies between the reactors; reactors 1 and 2 exhibited a more uniform community structure, with four genera making up over 70% of the total composition. In contrast, reactor 4 displayed more diversity, with seven genera making up 70% of the community.

Reactor 1 was dominated by methanogen *Methanobrevibacter arboriphilus* (35%) and *Eubacterium limosum* (22%) which together accounted for more than half of the community. *Eubacterium limosum*, a known acetogen, was previously detected in MES systems [39,61] and is capable of fixing CO_2_ into acetate via the Wood-Ljungdahl Pathway (WLP). Another key acetogen, *Sporomusa sphaeroides* [63], was present at 9% in reactor 1. In reactor 2, *M. arboriphilus* remained dominant (43%), followed by a *Propionicimonas* sp. (15%) and *E. limosum* (10%). Reactor 4 exhibited a distinct community composition with a departure from *M. arboriphilus* abundance (4%) and complete absence of *S. sphaeroides*. Here, *E. limosum* and *Clostridium aromativorans* constitute 40% of the total abundance. *C. aromativorans* is a recently identified species isolated from the pit mud of a fermentation pit in China [64]. *C. aromativorans* is closely related to *Clostridium luticellarii* FW431 (97.42% 16S rRNA gene sequence similarity) and has not yet been reported in chain elongation or gas-fermenting systems, with only putatively inferred butyrate-producing capabilities. It belongs to the Clostridia class, a class that plays a crucial role in chain elongation during gas fermentation [65,66].

### High quality genomes were constructed from metagenomic data

Across the three reactors, seven to nine high-quality metagenome-assembled genomes (MAGs) were identified, with contig counts ranging from 1 to 60 per bin. Generally, more fragmented bins corresponded to lower relative abundance, except in reactor 4, where the third most abundant bin comprised 56 contigs. Six circular genomes were successfully assembled across the three reactor metagenomes, including those of *C. aromativorans* and *S. sphaeroides*, underscoring the high quality of the sequencing data and providing insight into genome architecture (**Supplemental information 1; Tables S2 & S3**).

A genome blast indicated all species described above to be phylogenetically consistent across the different reactors, with a 95% similarity threshold **(Figure 2E and Supplemental information 2**). MAGs that were taxonomically identified up until species level were aligned to their most closely related NCBI reference genome given by GTDBTK with similarity threshold of 98% (**Supplemental information 3**). The alignment of the most abundant species in the metagenome to their reference, indicated the MAGs wear complete. The recovered genome of *Clostridium aromativorans* consisted of one circular fragment and provides a more complete and contiguous representation of the genome than the current reference [64] which is fragmented into more than 70 contigs **(Figure 2F**).

### Functional characterisation of MES community with meta-proteomics indicates expression of carbon fixation, chain elongation and methanogenesis pathways

Metaproteomic data allows us to further examine the functional architecture and role of each species in the communities. Relative proteome abundances indicate *Eubacterium*, *Sporomusa* and *Methanobrevibacter* as the most abundant genera in reactors 1 and 2, making up over 50% of the community. In reactor 4, over 40% of the community is made up by *Eubacterium* and *Clostridium* (**Figure 2D)**. This corresponds well with the metagenomic data. However, some key differences are present. Metaproteomic analysis detected a factor 3-4 times lower amount of *Methanobrevibacter* across all three reactors when compared to the metagenomic data. Similarly, metaproteomic data detected the consistent presence of *Propionicimonas* in each reactor, however with a two-fold lower abundance. Conversely, the metaproteomic data indicated a factor 2-3 higher abundance of *Sporomusa* in reactors 1 and 2.

A manual search of enzymes involved in five different carbon fixation pathways towards acetate, ethanol or lactate was performed to establish the carbon fixation capacities of the microbial communities [67] **(Figure 3A, search terms in Supplemental information 1 Table S8**). More than 7% of the relative proteome abundance in each reactor is involved in carbon fixation. Similar functional analysis was done for two chain elongation pathways and two methanogenesis pathways, accounting both for more than 2% of the proteome abundance in each reactor, respectively **(Figure 3A)** [68,69]. Both acetoclastic and hydrogenotrophic methanogenesis pathways were observed in the metaproteomes. Proteins of nearly all steps of both pathways were identified within the *Methanobrevibacter* species, with the hydrogenotrophic pathway fully reconstructed in reactor 1 and 4, and the acetoclastic pathway in reactor 4 (**Supplemental information 1 Figures S14 & S15**). The methanogenic proteome is lowest in R4, which corresponds to the lower abundance of methanogenic archaea in this reactor.

**Figure 3.**
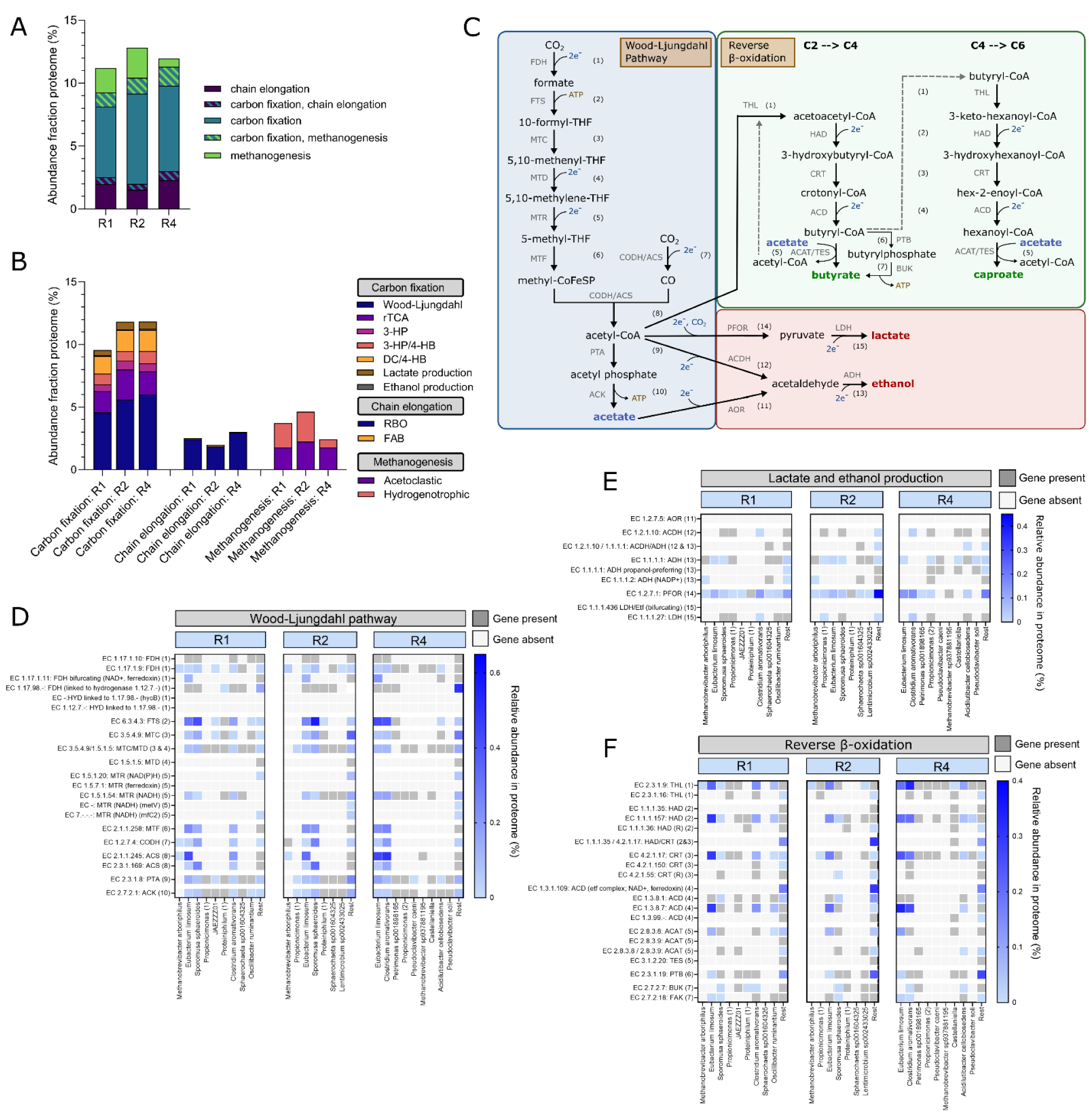
Functional analysis of the metaproteome isolated from three microbial electrosynthesis (MES) reactors. A: The abundance fraction in the metaproteome of (I) carbon fixation enzymes to acetate, ethanol or lactate production, (II) chain elongation of acetate to butyrate and caproate and (III) methanogenesis. B: Relative abundance of proteins in metaproteome of three categories (I) different carbon fixation routes to acetate, ethanol (from acetyl-CoA) or lactate (from acetyl-CoA), (II) chain elongation and (III) methanogenesis. The search terms that were used to classify the annotated proteins in each category and pathway were obtained from the KEGG database and can be found in supplemental information 1 table S8 [73]. The Wood-Ljungdahl Pathway (WLP) also includes the metaproteome fraction of the glycine cleavage complex (alternative route in WLP). Some pathways have overlapping EC and KO search terms and are therefore accounted for in the fraction of both pathways, making it difficult to determine the exact role of the protein. An example of such ambiguous KO numbers is the CODH/ACS complex involved in both WLP and acetoclastic methanogenesis. This might result in an overestimation of the abundance of some pathways. The overlapping fractions due to these ambiguities are detailed in supplemental information 4. C: Overview of putative MES product formation pathways of acetate, butyrate, caproate, ethanol and lactate. The overview includes CO2 fixation into acetate via the Wood-Ljungdahl Pathway (WLP), chain elongation to butyrate and caproate via the reverse beta-oxidation (RBO) and ethanol and lactate production from acetyl-CoA. D,E,F: Heatmaps showing the gene presence and protein abundance of the individual steps of the WLP (D), ethanol and lactate production from acetyl-CoA (E) and RBO (F) per MAG in all three reactors. Gene presence is shown in dark grey and protein abundance (%) in light to dark blue gradient.

### The Wood-Ljungdahl pathway as the most predominant carbon fixation route

The Wood-Ljungdahl Pathway (WLP) is known as the primary route of carbon fixation in most acetogens and is particularly relevant for MES [70,71]. Metagenomic analysis confirms the presence of the complete WLP in the microbial community **(Figure 3B)**. Further evidence of its metabolic role is provided by metaproteomic data which reveals that proteins associated with the WLP account for more than 4% of the total relative protein abundance across three reactors. This suggests that the WLP is the predominant carbon fixation pathway in these systems. Given it is the most energetically efficient pathway, it is unsurprising that the WLP is the primary mechanism for carbon fixation [72]. Additionally, proteins linked to alternative carbon fixation pathways, such as reverse tri-carboxylic acid cycle (rTCA) and the dicarboxylate-4-hydroxybutyrate cycle (DC/4HB cycle), each contribute only around 1-2% of the meta-proteome across the three reactors. For most steps in these two pathways, corresponding genes were identified in the metagenomic data; however, in contrast to the WLP, no single MAG revealed the complete set of any of the other carbon fixation routes that were investigated (**Supplemental information 1 Figures S16-S19**). The absence of their full complement of genes in a single genome and lower proteome abundance suggests a limited role for these alternative carbon fixation routes in this system.

### E. limosum, C. aromativorans and S. sphaeroides key players in carbon fixation

To identify key species involved in acetate production via WLP, we examined the presence of WLP-associated genes in MAGs **(Figure 3C & 3D**). Our analysis revealed the complete WLP being present in *Eubacterium limosum, Clostridium aromativorans*, and *Sporomusa sphaeroides*. Metaproteomic data further confirmed the presence of these proteins, as all WLP-associated proteins were detected withing these species. We assessed the contribution of each MAG to the WLP-related proteins. In reactor 1, *Eubacterium limosum* accounted for 37% of the WLP-related proteins, followed by *Sporomusa sphaeroides* with 33% and *Clostridium aromativorans* with 20%. In reactor 2, *Sporomusa sphaeroides* and *Eubacterium limosum* remained the dominant contributors with 37% and 25%, respectively. The residual WLP proteome mainly belongs to *Clostridium aromativorans* and an unknown bacterium, which both did not pass the checkM quality test and/or the abundance threshold of >2% in this reactor. In reactor 4, *Eubacterium limosum* and *Clostridium aromativorans* showed the highest relative WLP-related protein abundances at 36% and 37%, respectively. Overall, across the three reactors, the three key species contributing to carbon fixation via the WLP were consistently *Eubacterium limosum, Sporomusa sphaeroides* and *Clostridium aromativorans*. This aligns with the metagenomic analysis, which identified these species as among the most abundant members of the microbial community.

### Butyrate and caproate production is associated with reverse β – oxidation pathway

The two pathways analysed for chain elongation of acetate into butyrate and caproate are the reversed β-oxidation (RBO) and the fatty acid biosynthesis (FAB). While both pathways may result in production of short to medium chain carboxylic acids, the RBO seems to play an important role in other MCCA-producing processes, such as anaerobic fermentation from acetate using ethanol or lactate as electron donor [74,75] and has been the hypothesized chain elongation process in another MES system [43]. This pathway enables the sequential elongation of an acyl-CoA molecule with two carbons in a cyclic process. Metagenomic analysis confirms the presence of all genes required for both RBO and FAB pathways, while metaproteomic data provides further evidence of their expression **(Supplemental information 1 Figure S20)**. Based on relative abundance of these pathways in the proteome of each reactor, RBO is the dominant route with at least a ten-fold higher abundance compared to FAB (**Figure 3B**). This suggests that RBO is the primary mechanism for butyrate and caproate production across all three reactors.

### *E. limosum* and *C. aromativorans* as key contributors to butyrate and caproate production

Similar to carbon fixation, the presence of genes encoding the RBO pathway was investigated via metagenomics, and expression of RBO-related proteins was confirmed via meta-proteomics. While the full putative RBO pathway was identified in nearly all MAGs across the three reactors, complete expression of the pathway was limited to *Eubacterium limosum* and *Acidilutibacter cellobiosedens,* as indicated by metaproteomic data. Notably, expression of a near-complete RBO pathway was observed in multiple species, but proteins responsible for the final enzymatic conversion of acyl-CoA to carboxylic acid were consistently not detected (or absent) **(Figure 3C & 3F**). If the presence of a near-complete pathway indicates a species’ contribution to this metabolic function, the key players involved in chain elongation via the RBO pathway were identified based on the relative abundance of RBO-related proteins per MAG.

In reactor 1, most RBO-related proteins were attributed to *Eubacterium limosum* with 49%, followed by *Clostridium aromativorans* with 18%. Additionally, 23% of RBO-related proteins were found in the residual fraction, consisting of unbinned contigs and bins which did not pass the quality control. Similarly, in reactor 2 the highest number of relative abundance of RBO-related proteins belong to *Eubacterium limosum* with 41%. Most of the rest of the detected RBO proteins are found in the residual fraction, which, in this reactor, includes *Clostridium aromativorans.* In reactor 4, *Eubacterium limosum* again seems to be the highest contributor to the RBO pathway, with 38% of RBO-related proteins belonging to this MAG, closely followed by *Clostridium aromativorans*, contributing 33% of RBO-related proteins. This reactor contained *Acidilutibacter cellobiosedens,* with evidence of the complete RBO pathway, however, with a relative abundance of 7%, this species could not be identified as a key contributor to chain elongation via the RBO pathway in this community.

### Production of ethanol and lactate as potential electron mediators for chain elongation

As it was shown that chain elongation most likely occurs through the RBO pathway, question remains on what the electron carrier might be for production of butyrate and caproate in these MES systems. Ethanol and lactate are products of interest and potential electron carriers for chain elongation to MCCAs [76]. We explored the presence of genes and proteins in the metagenomic and -proteomic data connecting either acetyl-CoA or acetate to ethanol through acetaldehyde and acetyl-CoA to lactate via pyruvate (**Figure 3C)**. In all three reactors, the total relative abundance of proteins involved in ethanol and lactate production in the metaproteome was at least 0.05% and 0.42%, respectively (**Figure 3B**). We identified presence of acetaldehyde dehydrogenase and alcohol dehydrogenase encoding genes in the genome of 9 MAGs. However, expression of both enzymes was only detected in *Aciludibacter cellobsediens* in reactor 4 and the rest fraction in reactor 2 and 4. Genes encoding enzymes for lactate production from acetyl-CoA were identified in 9 MAGs, whilst evidence for protein expression was only observed in *Clostridium aromativorans* in reactor 1 and 4 (**Figure 3E**). In none of the MAGs, the NAD+/ferredoxin bifurcating lactate dehydrogenase was identified.

### Hypothetical cross-feeding interactions between community members

In microbial communities, the functional contribution of individual species critically depends on interactions such as cross-feeding of secondary metabolites. For rational optimisation of MES systems, it is important to understand how community members depend on each other for growth or product formation. Since a minimal catholyte medium lacking vitamins and amino acids was fed to the MES reactors, we hypothesize that the viability of some members of the community might be sustained by cross-feeding interactions. However, the lack of experimental tractability of these communities (difficult accessibility of the biofilms within the reactors, presence of uncultured species) makes it challenging to determine potential metabolic interactions in the lab. A computational alternative that is widely used as a workhorse to determine metabolic capabilities from microbial genomes are genome-scale metabolic models (GEMs) [77]. However, in our system, building high quality GEMs is also hindered by our inability to obtain experimental data for single species (some of them yet uncultured) that is required to curate the models. One possibility to bypass this issue is to use statistical approaches to maximize the functional information from incomplete or uncurated metabolic networks [78,79]. Here we assessed the potential for cross-feeding of specific metabolites using the recently devised “metabolic network percolation algorithm” [78], a probabilistic model that assess the likelihood that a model is able to produce a given metabolite based on its available metabolic network (a “producibility metric”, PM). Briefly, this approach repeatedly determines whether a model is able to produce a metabolite in randomly varying intracellular metabolic environments, effectively conducting a Monte-Carlo gapfilling of the metabolic network (**Figure 4A**). We focused specifically on B vitamins and aminoacids, as these are commonly found auxotrophies and are often expensive components that need to be added to the medium in similar systems (e.g. syngas fermentation, [80,81]) (**amino acid PMs; Supplemental information 1 Figure S22**). We started by reconstructing the metabolic networks with CarveMe [82] using either only genomic data, or additionally restricting it based on proteomic evidence. We next calculated the PM value for 9 B vitamins and 21 amino acids, and for 25 species models in all three microbial communities. We additionally included for each community a “mixed bag” model containing the rest fraction, i.e. all reactions that could not be assigned to a single species (Methods). We considered that a potential for cross-feeding of molecule *i* exists between two coexisting species *a* and *b* when the difference in PM, calculated from the genomic metabolic networks (**Figure 4A**), was larger than 0.5, i.e. |PM ^a^– PM ^b^|>0.5.

**Figure 4.**
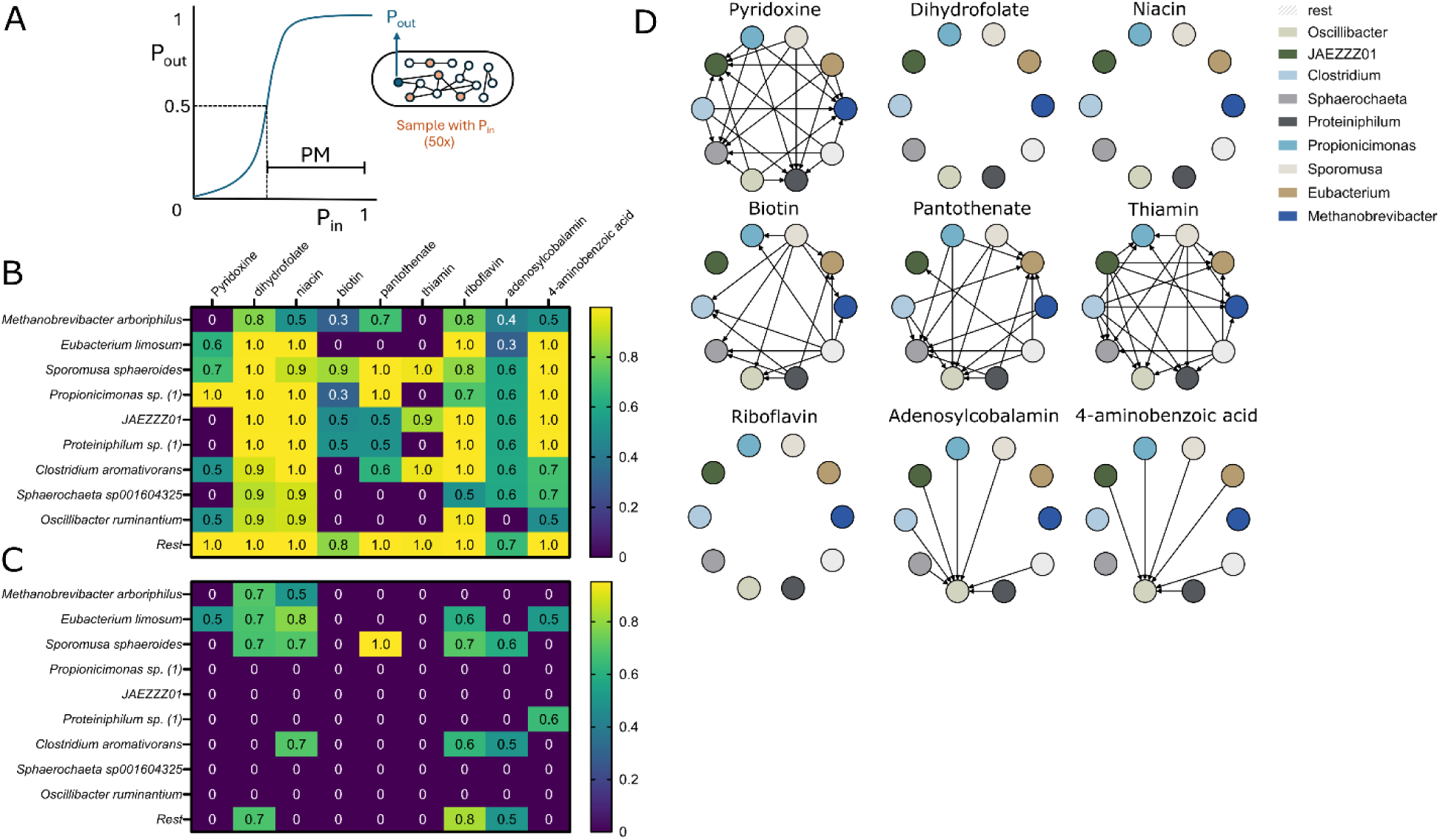
Vitamin B production capacities of the metabolic networks constructed from the isolated MAGs in reactor 1 to determine hypothetical cross-feeding interactions between community members. This reactor includes the three key role species, Eubacterium limosum, Clostridium aromativorans and Sporomusa sphaeroides. A: Figure adapted from Bernstein et al., a PM value close to 1 means high production robustness of that specific metabolite, a PM = 0 means the metabolite cannot be produced from any other metabolite in the metabolic network [78]. B: The producibility metrics (PM) calculated by the metabolic network percolation algorithm of Bernstein et al., 2019 with metabolic networks constructed from the isolated genomes in reactor 1 for several B vitamins (pyridoxine, dihydrofolate, niacin, biotin, pantothenate, thiamin, riboflavin, adenosylcobalamin and 4-aminobenzoic acid) [78]. C: The PM values calculated with the proteome carved metabolic networks of the isolated genomes of reactor 1 for the B vitamins. PM values of isolated genomes of reactor 2 and 4 are shown in supplemental information 1 figure S11. D: A schematic representation of potential cross-feeding interactions in the microbial community of reactor 1 based on the PM calculations shown in B. The nodes represent the different species and edges are drawn between species that have > 0.5 difference (|PM_i_^a^– PM_i_^b^|>0.5) in their PM value calculated from their genomic metabolic networks (arrow direction from species with high PM value to low).

Most species across the three communities have a PM > 0.6 for adenosylcobalamin, a form of vitamin B_12_, and a PM > 0.8 for dihydrofolate, a precursor of tetrahydrofolate, which are both important cofactors of the WLP [70,71,83] (**Figure 4B**; **reactor 1. Supplemental information 1 Figure S21; reactor 2 and 4**), suggesting the production of these is widespread. However, we identified significant potential for cross-feeding of pyridoxine (B_6_), pantothenate (B_5_), biotin (B_7_) and thiamin (B_1_) based on their differences in PM values between specific members of the communities **(Figure ^4^D).** High abundant species such as *S. sphaeroides* and *C. aromativorans* have PM values > 0.5 for these vitamins (except for biotin in *C. aromativorans*), whilst *E. limosum* is potentially dependent on exchange of pantothenate, thiamin and biotin by other members. As the producibility capacities are determined in randomly sampled environments, this cannot directly be linked to the metabolite production in MES conditions. Therefore, we also determined producibility capacities of MES context specific metabolic networks by carving them with the proteome. Considering the proteome carved network producibility of the B-vitamins, *S. sphaeroides, E. limosum* and *C. aromativorans* are hypothesized to be the main suppliers of most vitamins to the community in which they are present (**Figure 4C; reactor 1, Supplemental information 1 Figure S21; reactor 2 and 4**). Exceptions are biotin and thiamin which have a producibility of 0 in all three metaproteomes, as well as 4-aminobenzoic acid (B_10_) in reactor 2 and dihydrofolate in reactor 4. Overall, our data suggests that cross-feeding B vitamins is important for sustaining the structure of the communities.

## Discussion

In this work, we applied a unique strategy combining high resolution multi-omics with metabolic modeling to elucidate the structure and function of three high performing MES biofilm communities. Our metagenomic pipeline allowed us to reconstruct high quality genomes of 25 MAGs present in our reactors, including 6 fully circular genomes, and the most complete genome reconstruction of *Clostridium aromativorans* reported to date. Functional analysis revealed *E. limosum*, *S. sphaeroides* and *C. aromativorans* as key members of the communities, with the ability to both fix CO_2_ through the Wood-Ljungdahl and perform chain elongation through reverse β-oxidation. In particular, we report for the first time the presence of *C. aromativorans* in gas-fermenting and chain elongation systems, as well as its ability to produce C4-C6 MCCAs. We also report for the first time the capability of *E. limosum* to produce C4-C6 from CO_2_ and electricity. The presence of proteins for all steps in the transformation of acetyl-CoA to lactate or ethanol in the proteome might suggest these compounds could act as electron carrier intermediates [76]. Lactate production in MES has not been reported to date. Finally, meta-genome-scale metabolic modeling analysis suggests the communities might be sustained by the cross-feeding of specific cofactors (pyridoxine, pantothenate, biotin and thiamin). Altogether, our results qualitatively expand previous characterizations of mixed microbial communities in this system, paving the way for their rational engineering.

### High quality MAGs were constructed from the isolated metagenomes

The obtention of six fully circular genomes from the metagenomes indicates a high-quality sequencing data, achieved using a strategy of polishing long-read metagenomes with short reads. The metagenome of reactor 4 is however more fragmented than the metagenomes of reactor 1 and 2, possibly due to storage and freeze-thaw cycles causing DNA deterioration [84], as the DNA of reactor 4 was processed last. Additionally, differences in flow cell integrity may have impacted sequencing output quality [85]. Discrepancies between metagenomic and -proteomic relative abundances can be introduced by several factors, including the extraction procedures employed by the different methods as well as overexpression, cell volume – and viability fraction [86]. Regardless of these differences, the relative abundance profiles of the top taxonomies as indicated by the two meta-omics analyses were consistent. In comparison, 16S amplicon sequencing of the communities differed more substantially, indicating that our multi-omics approach was able to overcome the limitations and biases of the more commonly used 16S sequencing approach for estimating relative abundances [87].

### Lower abundance of methanogenic species observed in highest performing reactor

The highest butyrate and caproate production rates were found in the reactor with the highest abundance of *C. aromativorans*, suggesting a central role of this recently identified species in product formation. An interesting pattern is also seen in the abundance of methanogens. In particular, reactors 1 and 2 showed a high abundance of methanogens (*Methanobrevibacter* genus), which likely contributed negatively to desired product formation by introducing substrate competition for CO_2_ and H_2_. Additionally, methanogens can capture the acetate produced by acetogens, increasing methane production in detriment of valuable MCCAs [88]. Inhibiting methanogenesis with sodium 2-bromoethanesulfonate (BESA) has been attempted, but its effectiveness remains limited, possibly due to microbial adaptation or BESA metabolism by other anaerobic bacteria [89]. In contrast, the lower abundance of methanogenic archaea in reactor 4 is a potential reason for the higher performance in product formation of this community. Our data also revealed the presence of two species from the *Propionicimonas* genus, which had not been previously associated with MES systems, but had been detected in microbial fuel cells hypothetically contributing to extracellular electron transfer [90]. The function of *Propionicimonas* in MES biofilms must be investigated further. More broadly, relating MES reactor product profiles to the composition of their microbial communities would be immensely valuable. The challenge of this approach lies in our currently limited ability to run a large number of replicate MES reactors. In the future, the influence of specific species and their abundance within a mixed community could be also investigated using synthetic consortia. This approach could enable a tighter control over these biological systems and could pave the way to higher product specificity.

### Functional duality of *E. limosum, C. aromativorans* and *S. sphaeroides* in their carbon fixation and carbon chain elongation abilities

Interestingly, the gene presence of all steps in WLP and RBO plus proteomic evidence of most steps in these pathways, in *E. limosum, C. aromativorans* and *S. sphaeroides* suggests that these species play a dual role in carbon fixation and chain elongation. While butyrate and caproate production by *E. limosum* has been previously reported, it has not been observed under conditions where CO_2_ and electrons from an electrode and/or H_2_ serve as the sole substrates [91,92]. Energetically, butyrate production from CO is feasible, yielding 3.5 mol ATP/mol butyrate in *E. limosum* KIST621. However, production from H_2_ and CO_2_ lacks sufficient reducing equivalents in the absence of additional substrates [93]. The closest sequenced strain of *E. limosum* to the strain in our reactors (ATCC 8486, **Supplemental information 3)**, has been reportedly unable to produce butyrate even from CO [94]. From the thermodynamic standpoint, previous calculations showed that chain elongation from acetate from H_2_ is infeasible [76]. This raises the question of what mechanisms are making chain elongation possible in our system. An interesting hypothesis is the possibility of spatial segregation within the biofilm, with *E. limosum* near the cathode core primarily performing acetogenesis, while those at the biofilm periphery engaging in chain elongation. This scenario would require cross-feeding of short-chain fatty acids across the biofilm (e.g. acetate away from the cathode) or reduced electron carriers such as ethanol or lactate. A similar metabolic duality was observed for *C. aromativorans*. Genome alignment of *C. aromativorans* revealed close similarity to *Clostridium sp*. BL-3, a strain capable of producing caproate from lactate and isolated from a chain elongating reactor [95]. Although neither lactate nor ethanol production have been measured experimentally in our MES reactors, their detection might be difficult if they fail to accumulate due to their rapid consumption [66]. Such a scenario is partially supported by our metagenomic and metaproteomic data showing the presence of genes and proteins involved in ethanol and lactate production from acetyl-CoA. However, the expression of these enzymes was detected in a small fraction of MAGs, suggesting that this capability may be restricted to a limited set of members of the community, or may be dependent on cross-feeding of other intermediates (e.g. acetaldehyde or pyruvate). Aside from this, lactate production from H_2_ and CO_2_ would only be energetically possible in the presence a bifurcating methylene-THF reductase [96]. Further work is needed to elucidate the precise metabolic interactions, most critically regarding possible electron donors, governing chain elongation in these reactors.

### High producibility capacities for B vitamins in *E. limosum, C. aromativorans* and *S. sphaeroides*

A critical aspiration in the field of microbial electrosynthesis is to move from undefined, mixed communities to defined synthetic communities amenable to rational manipulation and engineering. To achieve this goal, it is critical to uncover the potential metabolic dependencies sustaining community structure and function. We employed here a cutting-edge modeling methodology, metabolic network percolation, designed to uncover potential interactions using partially complete metabolic network reconstructions [78]. Based on the calculated B vitamins (and amino acids) producibility capacities of the high abundant species *E. limosum, C. aromativorans* and *S. sphaeroides,* these would opt for good candidates when moving towards defined consortia in MES using minimal media. The potential auxotrophy of *E. limosum* for pantothenate and biotin has been shown experimentally before in other *E. limosum* strains [97], but is less clear in the case of thiamine. However, vitamin auxotrophy could be circumvented in defined consortia through cross-feeding with other members. Although the metabolic network percolation method extracts maximal information from the available data, the limitations of having incomplete draft metabolic network reconstructions (e.g. because of lack of gene annotation or detection limits in the proteome) should be kept in mind when drawing conclusions. The quality of the model reconstruction pipeline may be improved by including the data from other annotation sources, such as the KO numbers identified by eggNOG-mapper.

### Concluding remarks

Our cross-disciplinary strategy allowed us to reconstruct multiple high-quality genomes, unveil their metabolic capabilities and the ecological interactions potentially sustaining community structure. The integration of high-resolution multi-omics with advanced metabolic modeling techniques represents an emerging trend in the broader field of microbial communities and microbiome science [98,99]. This strategy allowed us to identify a potential dual function in three of the dominant species in our reactors, as *Eubacterium limosum, Sporomusa sphaeroides* and *Clostridium aromativorans* are all capable of both CO_2_ fixation and chain elongation. The latter species was previously not reported in MES communities, broadening the portfolio of potentially interesting species for this application. Our in-depth characterization of MES biofilm communities paves the way towards the rational engineering of high-performing electrosynthetic communities. An exciting future perspective is the construction of defined synthetic communities (SynCom), which might be more desirable in the industrial application of MES. Our study provides crucial guidance for future efforts in strain isolation, media design, and testing of defined consortia that will be essential to achieve a feasible technology. In summary, our work represents a foundational piece in the development of a future technology that harnesses microbial communities for fixation of CO_2_ into valuable products.

## Materials and Methods

### MES reactor operation and sampling

This work describes the microbial characterization of three Microbial Electrosynthesis (MES) reactors previously described by Winkelhorst *et. al.* These reactors were set up and conditionally maintained as replicates, identical time and source of inoculation, reactor design, temperature, pH, catholyte, and flow-rate [58]. Biomass harvesting was performed at the end of the 194 day run by excising cross-section samples of the carbon-felt cathode material, which were subsequently stored at −80 °C in 50:50 catholyte:glycerol.

### DNA extraction

To extract DNA from the biofilm samples in the MES reactors for both 16S rRNA sequencing and whole genome sequencing, the DNA extraction was performed as described previously in Cabau-Peinado et al., 2024 [39].

### 16S rRNA illumina sequencing

The extracted DNA samples were sent to Novogene (UK) for 16S rRNA sequencing. The microbial analysis via 16S rRNA sequencing was performed following the microbial community analysis method previously described in Cabau-Peinado et al., 2024 [39].

### Metagenome sequencing

Whole genome sequencing was performed using long-read sequencing from Nanopore technology. Library preparation was conducted using the SQK-LSK112 ligation sequencing kit (Oxford Nanopore Technologies, Oxford, UK) following the manufacturer’s protocol. Sequencing was performed in R10.4 flow cells, docked into MinION devices, using MinKNOW software (version 22.05.5) (Oxford Nanopore Technologies, Oxford, UK).

An additional short-read sequencing step was performed to polish the Nanopore sequences. Samples from the three reactors were sequenced at Macrogen with TruSeq DNA PCR-Free library prep kit. Sequencing was performed on Illumina NovaSeq platform to generate 151 bp paired-end reads with total sequences of ∼10 gigabases for each reactor.

### Metagenome data processing

ONT Guppy basecalling algorithm (version 6.3.4) (https://nanoporetech.com/software/other/guppy) as applied on the raw signal data files (FAST5) from the sequence runs, yielding total sequences of 5.59 gigabases (R1), 9.05 gigabases (R2) and 5.23 gigabases (read length >= 1000 bases). With a N50 read length of 3719 bases (R1), 4561 bases (R2) and 3700 bases (R4). Mean quality score of the three reactors were 13.5 (R1), 14.1 (R2) and 12.8 (R4).

Reads of length 1000 base pairs and larger were assembled into contigs using Flye assembler (version 2.9.1-b1780 with option --meta) [100]. Illumina reads were mapped to the assembled contigs using BWA (version 0.7.18-r1243-dirty) [101]. The resulting SAM file was sorted into a BAM file using Samtools (version 1.18) [102]. After mapping, the nanopore assembled contigs were polished with the Illumina reads using Pilon (version 1.18) [103]. Illumina reads were again mapped to the polished contigs by BWA and sorted by Samtools for binning. Three different binning tools with the following non-default settings were applied; I) concoct, II) metaBat2 with minimal contig length and depth of 1000bp and 2 respectively and III) maxbin2 [104–106]. The results of the three binning tools were consolidated to the final bins and an unbinned fraction using DASTOOL with score threshold of 0.6 (version 1.1.6) [107].

The quality of the bins was analysed with checkM2 [108] and filtered based on a completeness higher than 95% and contamination lower than 5%, resulting in the metagenome-assembled genomes (MAGs) discussed in this study **(Supplemental Information 1 Table S4)**. Sequences not meeting these criteria as well as MAGs having a relative abundance lower than 2% and unbinned contigs were grouped under “rest”. The MAGs were taxonomically classified with GTDBTK 2.4.0 [109]. To further asses the completeness and the quality of the isolated genomes, the MAGs that were taxonomically identified up until species level were aligned with BLASTn [110] to their closest genome reference assigned by GTDBTK, which were obtained from the NCBI RefSeq database [111]. The BLASTn hits were filtered based on an e-value < 1e-50 and a percentage identity > 98%. Taxonomically identical MAGs in the three communities were aligned using the BLASTn tool, with e-value < 1e-50 and percentage identity > 95%, to determine species similarity in the reactors. Circos was used to construct plots for alignment visualisation [112]. Samtools was used to determine the coverage of each MAG based on the mapped Illumina reads, to calculate the relative abundance of each MAG in the communities according to the following equation:

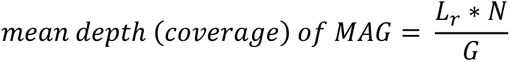

with L = length of read, N = number of reads and G = MAG length [113]. Relative abundance was then calculated by dividing the mean depth of the MAG by the sum of the mean depth of all MAGs. Prokka (version 1.14.6) was used to identify and annotate coding sequences in the metagenomes [114]. Additionally, the with Prokka identified coding sequences in the metagenomes were used as input for eggNOG-mapper (version 2.1.12) to functionally annotate the metagenomes based on orthologous groups, including DIAMOND with –more-sensitive mode for taxonomic classification [115,116]. These annotated metagenomes were used as blueprint for proteomic analysis.

### Protein extraction and proteolytic digestion

Samples were taken from three locations at the cathode in each reactor. From there, protein extraction was performed using a modified protocol as described previously (den Ridder et al., 2023). Briefly, biofilm samples grown on carbon membrane material (approximately 150 mg) were mixed with 200 µL of 50 mM triethylammonium bicarbonate (TEAB) containing 1% sodium deoxycholate (NaDOC) and 200 µL of B-PER™ reagent (Thermo Fisher). Biofilm material and cells were disrupted by glass bead beating using acid-washed 150–212 µm glass beads (Sigma-Aldrich) for three cycles of 1.5 min beating, each followed by 30 s cooling on ice. Lysates were heated at 80 °C for 3 min, sonicated for 10 min, and centrifuged at 14,000 × g for 10 min at 4 °C to remove debris. Supernatants were collected in LoBind Eppendorf tubes for protein precipitation. Proteins were precipitated by adding trichloroacetic acid (TCA) at a 1:4 ratio and incubating for 30 min at 4 °C, followed by centrifugation at 14,000 × g for 15 min. Resulting protein pellets were washed with ice-cold acetone, briefly centrifuged, and redissolved in 6 M urea prepared in 200 mM ammonium bicarbonate (ABC). Reduction was carried out with 10 mM dithiothreitol for 1 h at 37 °C, followed by alkylation with 20 mM iodoacetamide for 30 min at room temperature in the dark. Samples were diluted with ABC to reduce the urea concentration below 1 M and digested overnight at 37 °C with sequencing-grade trypsin (Promega) at a 1:100 enzyme-to-protein ratio. Peptides were desalted using Oasis HLB µElution plates (Waters). The peptide fraction was loaded onto the cartridge under aqueous conditions, washed twice with 350 μL of 5% methanol in water, and eluted in two steps with 200 μL of 80% methanol containing 2% formic acid and 200 μL of 1 mM ABC in 80% methanol. The combined eluate was dried in a SpeedVac concentrator. Dried peptides were reconstituted in 3% acetonitrile with 0.01% trifluoroacetic acid, quantified using a NanoDrop spectrophotometer, and diluted to approximately 0.5 mg/mL for subsequent shotgun proteomic analysis.

### Shotgun metaproteomic analysis

Shotgun proteomic analysis was performed as described previously [118]. Briefly, from each sample, an aliquot corresponding to approximately 500 ng of protein digest was analysed in duplicate using an EASY-nLC 1200 nano-liquid chromatography system equipped with an Acclaim PepMap RSLC C18 column (50 μm × 150 mm, 2 μm; Thermo Fisher Scientific, Germany) coupled to a QE Plus Orbitrap mass spectrometer (Thermo Fisher Scientific, Germany). The flow rate was set to 350 nL/min using a linear gradient from 5% to 30% solvent B over 90 min, followed by 30% to 60% over 25 min, and re-equilibration to starting conditions. Data were acquired between 5 and 120 min. Solvent A was water containing 0.1% formic acid (FA), and solvent B was 80% acetonitrile in water with 0.1% FA. The Orbitrap was operated in data-dependent acquisition (DDA) mode, acquiring peptide signals from m/z 385–1250 at 70k resolution in full MS scans, with a maximum injection time (IT) of 100 ms and an automatic gain control (AGC) target of 3E6. The top 10 precursors were selected for fragmentation using higher-energy collisional dissociation (HCD). MS/MS spectra were acquired at 17.5k resolution with an AGC target of 2E5, IT of 75 ms, isolation width of 2.0 m/z, and normalized collision energy (NCE) of 28. Data processing of the metaproteomic samples was performed as described by [86]. Mass spectrometric raw data were searched using PEAKS Studio X (Bioinformatics Solutions Inc., Canada) against a database generated from the corresponding metagenomic sequencing data (see above), allowing a 20 ppm precursor and 0.02 Da fragment mass error, with oxidation and deamidation set as variable modifications. Identified proteins were annotated using eggNOG-mapper functional assignments and DIAMOND-derived taxonomic lineages.

### Pathway expression analysis in metaproteome

Metaproteomic data from the three sampling locations at the cathode were combined to eliminate sampling biases and to get the most representative image for every reactor. The number of spectra for each identified protein was corrected for the mass of the protein and multiplied by a large number as follows:

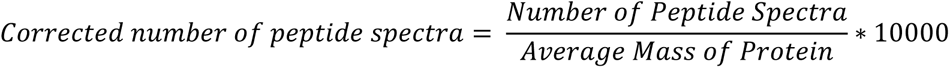

The corrected number of spectra for each protein was used to calculate the protein relative abundances. MAG relative abundance regarding protein expression was calculated by summing up the corrected spectral counts of all identified proteins of each individual MAG divided by the total amount of spectral counts in the proteome. Relative abundance of each individual protein was calculated by dividing the corrected number of spectra of the protein of interest by the sum of the corrected number of spectra of all identified proteins in the proteome. The detected proteins were traced back to the specific ORF accession numbers identified and assigned in the metagenomes by Prokka to analyse MAG specific pathway expression and function of the proteins. For functional analysis of the metagenomic and metaproteomic data, gene presence and protein expression of carbon fixation and chain elongation metabolic pathways, including Wood-Ljungdahl and reverse beta-oxidation were analysed based on the EC numbers and KO hits annotated by Prokka and eggNOG-mapper to the coding sequences. The KO identifier assigned by eggNOG-mapper was used for the manual search if available; if not, the EC number assigned by Prokka was used instead. An overview of the KO and EC search terms of the genes involved in the pathways of interest was obtained from the KEGG database [73] and is given in **Supplemental information 1 Table S8.** The relative metaproteome abundance of each MAG at the three separate locations can be found in **Supplemental information 1 Figure S2.** Functional analysis of the metaproteome at separate locations can be found in **Supplemental information 1 Figures S3-S13 and Tables S5-S7**. Python 3.10.13 and 3.11.4, Rstudio 2022.12.0 with R 4.4.1 and the readxl 1.4.3, tidyverse 2.0.0 and writexl 1.5.0 packages and Graphpad Prism 10.2.3 were used for data processing and visualization.

### Metabolic network construction and interaction analysis of the isolated MAGs

Hypothetical cross-feeding interactions between members in the MES communities were analysed with the metabolic network percolation algorithm of Bernstein et al., 2019 using draft metabolic networks of the individual MAGs constructed with CarveMe (version 1.6.2) [78,82]. Diamond was used with CarveMe for reaction annotation by aligning the by Prokka translated ORFs with default settings [115]. The bacterial universal model of CarveMe was used as template for the bacterial identified MAGs and the archaea universal model for the *Methanobrevibacter* isolated MAGs. The mixed-bag approach was used for construction of a metabolic model from the ‘rest’ group, containing bins that did not pass the quality and relative abundance constraints and the unbinned contig fraction [119,120]. Reactions for adenosylcobalamin and biotin were manually added to the draft metabolic models based on eggNOG-mapper annotation of the genomes as these reactions were missing from the CarveMe universal models. Metabolic reactions that were added to the models by CarveMe without genetic evidence were removed. Proteome carved metabolic networks of the MAGs were constructed by removing all metabolic reactions that were not detected in the metaproteome. A basic check for presence of free ATP generating fluxes in the metabolic models of the MAGs showed that none were present.

The metabolic network percolation algorithm enables the analysis of genome-scale models of uncultivated species for which curation of the metabolic model is difficult. A producibility metric (PM, between 0-1) for metabolites of interest was calculated to quantify the capabilities of producing the metabolite from the given network by randomly sampling the environment. A large PM indicates high production robustness of the specific metabolite given the metabolic network. A PM of 0 means the metabolite cannot be produced from any other metabolite in the network. The PMs of nine vitamin B metabolites (pyridoxine, dihydrofolate, niacin, biotin, pantothenate, thiamin, riboflavin, adenosylcobalamin, 4-aminobenzoic acid) - based on vitamin solution added in syngas fermentation [121], lipoate was excluded from the analysis as the universal CarveMe template had a PM of 0 for this metabolite - and of 21 amino acids (including selenocysteine) were calculated, sampling the intracellular metabolites, oxygen excluded at all times, in the draft metabolic networks. The following parameters were used in the algorithm; s = 1, samp = 50, noise = 0.3, n = 7, thresh = 0.01, runs = 10 and a = int_nt, the latter one meaning that the intracellular environment of the metabolic network is sampled. The PM average and standard deviation were calculated from the 10 runs (**Supplemental information 4**). Hypothetical interactions were determined by an arbitrarily chosen difference of 0.5 in metabolite PM values between members of the community. PM values were also determined for the universal prokaryotic template of CarveMe as a control, with and without the manually added reactions for adenosylcobalamin and biotin, and the metabolic networks constructed from the complete metaproteome in each reactor to determine the producibility capacity of the whole metaproteome (**Supplemental information 1 Figure S14**). MATLAB version R2021b was used for running the metabolic network percolation algorithm.

## Resource availability

### Data and code availability

The mass spectrometry proteomics raw data have been deposited in the ProteomeXchange consortium database with the dataset identifier PXD069469 and will be available at time of publishing. The whole metagenome sequencing data and assemblies will be available through the NCBI Bioproject repository (https://www.ncbi.nlm.nih.gov/bioproject) under the accession number PRJNA1375776. The developed python and R codes for processing the metadata and creating the figures will be available online at time of publishing.

## Acknowledgements

Co-funding was obtained from dsm-firmenich and from a supplementary grant ‘TKI- Toeslag’ for Topconsortia for Knowledge and Innovation (TKI’s) of the Netherlands Ministry of Economic Affairs and Climate Policy (CHEMIE.PGT.2021.003). MG, LJ, and DB want to acknowledge financial support from the Zero Emission Biotechnology programme of Delft University of Technology.

