## Supplemental Information 1 for "Linking structure to function in high performing electrosynthetic biofilm communities"

#### 1. Performance overview of reactors during last 20 days of operation

*Table S1. Overview of the performance of the last 20 days of operation of reactor 1 (R1), reactor 2 (R2) and reactor 4 (R4) from Winkelhorst et al., 2023.*

| | **Concentration (mmol/L)** | | | **Rate (mmol/L/day)** | | | **Faradaic efficiency** | | | **C2:C4:C6 ratio's** |
|---|---|---|---|---|---|---|---|---|---|---|
| | **C2** | **C4** | **C6** | **C2** | **C4** | **C6** | **C2** | **C4** | **C6** | **cmol** |
| R1 | 104 ±18 | 36 ±5 | 5,0±0,3 | 13±6 | 4±2 | 0,6±0,1 | 19%±6% | 16%±6% | 4%±1% | 7:5:1 |
| R2 | 95 ±15 | 12±4 | 0 | 14±2 | 2±1 | | 39%±10 % | 14%±4% | | 4:1 |
| R4 | 155 ± 11 | 97±1 1 | 7,0±0,5 | 20±4 | 12± 4 | 0,9±0,2 | 41%±8% | 64%±19 % | 7%±2% | 8:9:1 |

#### 2. Overview nanopore and Illumina sequencing analysis

*Table S2. Metadata of metagenomic sequencing results.*

| **Sample** | **# coding sequences identified by Prokka** | **# Hypothetical proteins** | **# contigs** | **# MAGs*** | **# circular genomes** |
|---|---|---|---|---|---|
| R1 | 117800 | 65544 | 6601 | 9 | 1 |
| R2 | 175078 | 72340 | 9563 | 7 | 4 |
| R4 | 176100 | 73958 | 17979 | 9 | 1 |

*Bins that passed the checkM2 quality test (completeness > 95% and contamination <5%) and have a relative abundance > 2%.

*Table S3. Circular genomes isolated from the metagenome of reactor 1 (R1), reactor 2 (R2) and reactor 4 (R4). Circular MAG with taxonomy Belilinea is has a relative abundance <2% and is included in the rest group.*

| **Sample** | **GTDBTK taxonomy** | **Length of circular contig (base pairs)** |
|---|---|---|
| R1 | *Propionicimonas sp.* | 2703843 |
| R2 | *Propionicimonas sp.* | 2703803 |
| R2 | *Sporomusa sphaeroides* | 4912660 |
| R2 | *Lentimicrobium sp002433025* | 4971150 |
| R2 | *Bellilinea sp.* | 4432006 |
| R4 | *Clostridium aromativorans* | 3814898 |

*Table S4. CheckM2 quality scores of all bins obtained from whole genome sequencing in samples of reactor 1 (R1), reactor 2 (R2) and reactor 4 (R4).*

| Sample | bin | GTDBTK taxonomy | checkM2 completeness | checkM2 contamination |
|---|---|---|---|---|
| R1 | concoct.35 | *Methanobrevibacter arboriphilus* | 100 % | 1 % |
| R1 | maxbin_out.002 | *Eubacterium limosum* | 100 % | 2 % |
| R1 | maxbin_out.003 | *Sporomusa sphaeroides* | 100 % | 5 % |
| R1 | maxbin_out.004 | *Propionicimonas (1)* | 100 % | 0 % |
| R1 | maxbin_out.005 | *JAEZZZ01* | 100 % | 0 % |
| R1 | metabat_bin.9 | *Proteiniphilum (1)* | 100 % | 1 % |
| R1 | maxbin_out.007 | *Clostridium aromativorans* | 100 % | 1 % |
| R1 | maxbin_out.008 | *Sphaerochaeta sp001604325* | 99 % | 2 % |
| R1 | metabat_bin.36 | *Proteiniphilum (2)* | 100 % | 10 % |
| R1 | metabat_bin.15 | *Oscillibacter ruminantium* | 100 % | 4 % |
| R1 | metabat_bin.37 | *Lentimicrobium sp002433025* | 96 % | 7 % |
| R1 | maxbin_out.012_sub | *Methanobrevibacter sp937881195* | 96 % | 3 % |
| R1 | metabat_bin.11 | *Oscillibacter* | 79 % | 7 % |
| R1 | concoct.50 | *Bilophila wadsworthia* | 94 % | 9 % |
| R1 | concoct.22_sub | *Oscillibacter sp023393155* | 89 % | 27 % |
| R1 | metabat_bin.31 | *Petrimonas sp002356435* | 77 % | 2 % |
| R1 | concoct.39 | *Sphaerochaeta associata* | 100 % | 10 % |
| R1 | concoct.7 | *JAQUCI01* | 82 % | 7 % |
| R1 | metabat_bin.17 | *JAAYUD01* | 82 % | 7 % |
| R1 | maxbin_out.026 | *S3-3* | 83 % | 12 % |
| R1 | maxbin_out.027_sub | *Petrimonas* | 87 % | 32 % |
| R1 | maxbin_out.028 | *Desulfovibrio sp900243745* | 57 % | 3 % |
| R1 | metabat_bin.39 | *x* | 59 % | 2 % |
| R2 | concoct.30 | *Methanobrevibacter arboriphilus* | 100 % | 0 % |
| R2 | concoct.31 | *Propionicimonas (1)* | 100 % | 0 % |
| R2 | maxbin_out.003 | *Eubacterium limosum* | 100 % | 2 % |
| R2 | concoct.26 | *Sporomusa sphaeroides* | 100 % | 5 % |
| R2 | maxbin_out.005 | *Proteiniphilum (1)* | 100 % | 1 % |
| R2 | concoct.2 | *Sphaerochaeta sp001604325* | 99 % | 2 % |
| R2 | concoct.11 | *Lentimicrobium sp002433025* | 97 % | 2 % |
| R2 | concoct.54 | *Oscillibacter* | 100 % | 2 % |
| R2 | maxbin_out.011 | *Bellilinea* | 99 % | 5 % |
| R2 | concoct.49 | *JAEZZZ01* | 100 % | 6 % |
| R2 | concoct.21 | *Azospira* | 100 % | 2 % |
| R2 | metabat_bin.51 | *JAQUCI01* | 96 % | 4 % |
| R2 | metabat_bin.39 | *Proteiniphilum (2)* | 86 % | 5 % |
| R2 | concoct.3 | *Acidilutibacter cellobiosedens* | 95 % | 1 % |
| R2 | concoct.19 | *Bilophila wadsworthia* | 100 % | 10 % |
| R2 | concoct.13 | *x* | 93 % | 7 % |
| R2 | maxbin_out.020 | *Sphaerochaeta associata* | 100 % | 18 % |
| R2 | metabat_bin.30_sub | *Oscillibacter* | 57 % | 8 % |
| R2 | concoct.39 | *Propionicimonas (2)* | 86 % | 5 % |
| R2 | metabat_bin.14 | *Oscillibacter sp023393155* | 94 % | 43 % |
| R2 | maxbin_out.023_sub | *Clostridium aromativorans* | 82 % | 8 % |
| R2 | maxbin_out.027 | *Intestinimonas massiliensis* | 93 % | 8 % |
| R2 | maxbin_out.035 | *JAAYUD01* | 76 % | 29 % |
| R2 | metabat_bin.25 | *Paludibacter sp029973975* | 65 % | 2 % |
| R2 | concoct.22_sub | *Seramator sp029975415* | 78 % | 2 % |
| R2 | concoct.36_sub | *Syner-03 sp002306075* | 63 % | 1 % |
| R2 | maxbin_out.029 | *UBA6107 sp012797695* | 66 % | 5 % |
| R4 | concoct.28 | *Eubacterium limosum* | 100 % | 2 % |
| R4 | maxbin_out.002 | *Clostridium aromativorans* | 100 % | 1 % |
| R4 | maxbin_out.004 | *Petrimonas sp001898165* | 99 % | 0 % |
| R4 | maxbin_out.003 | *Propionicimonas (2)* | 100 % | 0 % |
| R4 | concoct.14 | *Pseudoclavibacter caeni* | 100 % | 1 % |
| R4 | maxbin_out.007 | *Oscillibacter ruminantium* | 59 % | 11 % |
| R4 | metabat_bin.70 | *Methanobrevibacter sp937881195* | 95 % | 1 % |
| R4 | maxbin_out.009 | *Castellaniella* | 100 % | 1 % |
| R4 | concoct.34_sub | *Oscillibacter* | 100 % | 38 % |
| R4 | concoct.7 | *Acidilutibacter cellobiosedens* | 97 % | 3 % |
| R4 | metabat_bin.53 | *Proteiniphilum* | 83 % | 2 % |
| R4 | concoct.13_sub | *Methanobrevibacter arboriphilus* | 85 % | 5 % |
| R4 | metabat_bin.36_sub | *Pseudoclavibacter soli* | 96 % | 2 % |
| R4 | maxbin_out.014 | *x* | 93 % | 4 % |
| R4 | metabat_bin.71 | *Sphaerochaeta sp001604325* | 77 % | 6 % |
| R4 | concoct.21 | *S3-3* | 95 % | 9 % |
| R4 | concoct.49 | *Propionicimonas* | 86 % | 6 % |
| R4 | metabat_bin.51_sub | *Petrimonas sp002356435* | 82 % | 7 % |
| R4 | metabat_bin.68 | *Caproicibacter* | 89 % | 3 % |
| R4 | concoct.37_sub | *Achromobacter pulmonis* | 81 % | 8 % |
| R4 | metabat_bin.26_sub | *Rummeliibacillus suwonensis* | 67 % | 7 % |

#### 3. 16S, metagenomic and metaproteomic abundance

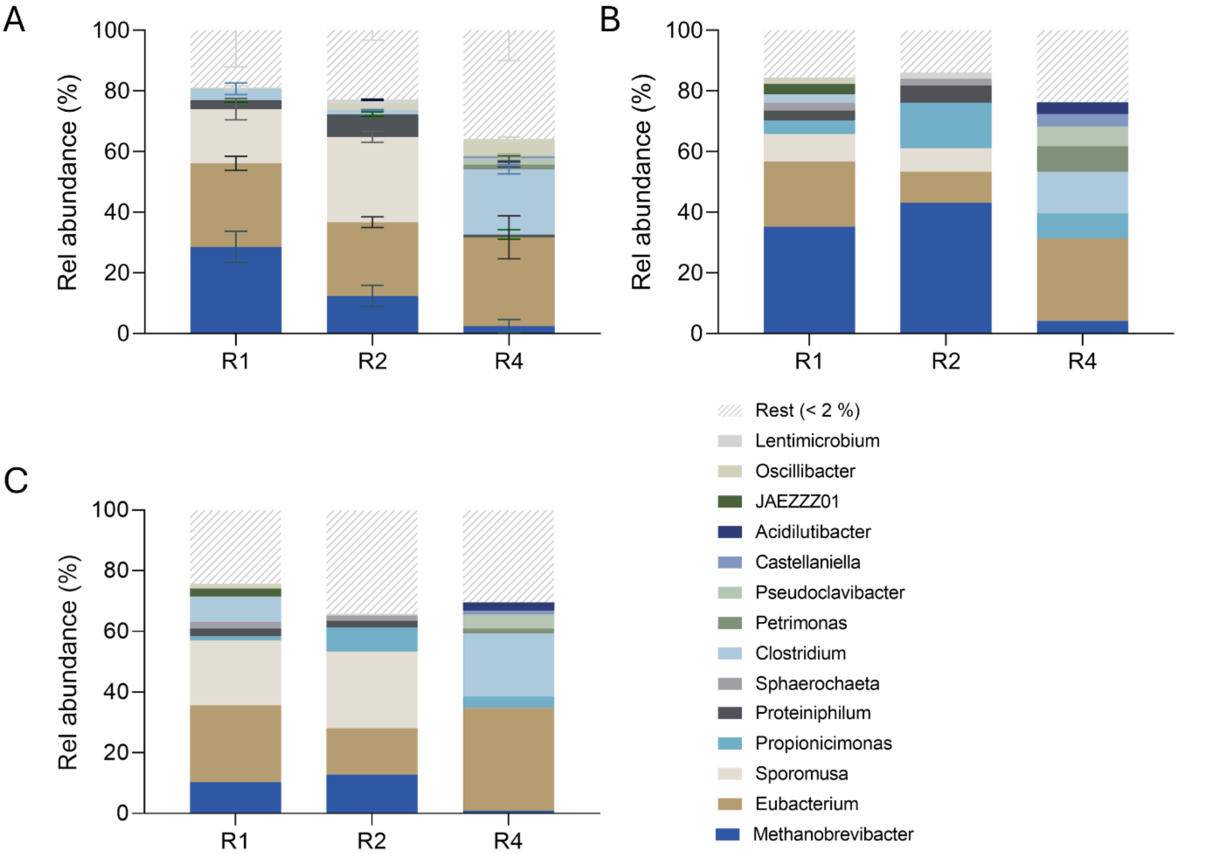


*Figure S1. Relative abundance of genera within the microbial communities of three MES reactors as according to 16S rRNA gene sequencing data (A), metagenomic sequencing data via nanopore, polished with Illumina short-read sequencing data (B), and metaproteomic data (C). "Rest" describes those sequences which did not meet the quality standards of CheckM2, were unbinned, or fell under a 2% relative abundance threshold. 16S rRNA gene sequencing was based on three technical replicates of DNA samples from reactors 1 (R1) and 4 (R4), and five technical replicates from reactor 2 (R2). Standard deviation between these replicates are included as error bars. Metagenomic and -proteomic data was based on single samples.*

We observed notable disparities between the results obtained from 16S rRNA gene sequencing and whole genome sequencing (WGS). Most notably, the genus of *Propionicimonas*, consistently detected present in all three reactors via WGS, was completely absent in 16S sequencing data. Additionally, WGS revealed a twofold higher relative abundance of *Methanobrevibacter* in reactors 1 and 2 when compared to 16S sequencing. The discrepancies may be attributed to biases introduced during amplification in 16S sequencing, including primer efficiency, specificity [1] and polymerase-induced errors. Additionally, variations in 16S rRNA gene copy number per genome can distort the relative abundance estimates. For example, *Sporomusa sphaeroides* contains twelve 16S rRNA gene copies, whereas *Methanobrevibacter arboripilus* possesses only two 16S rRNA gene copies, potentially skewing abundance calculations [2]. Despite correction tools, these biases remain unresolved.

#### 4. Proteomics abundance of MAGs at different locations samples

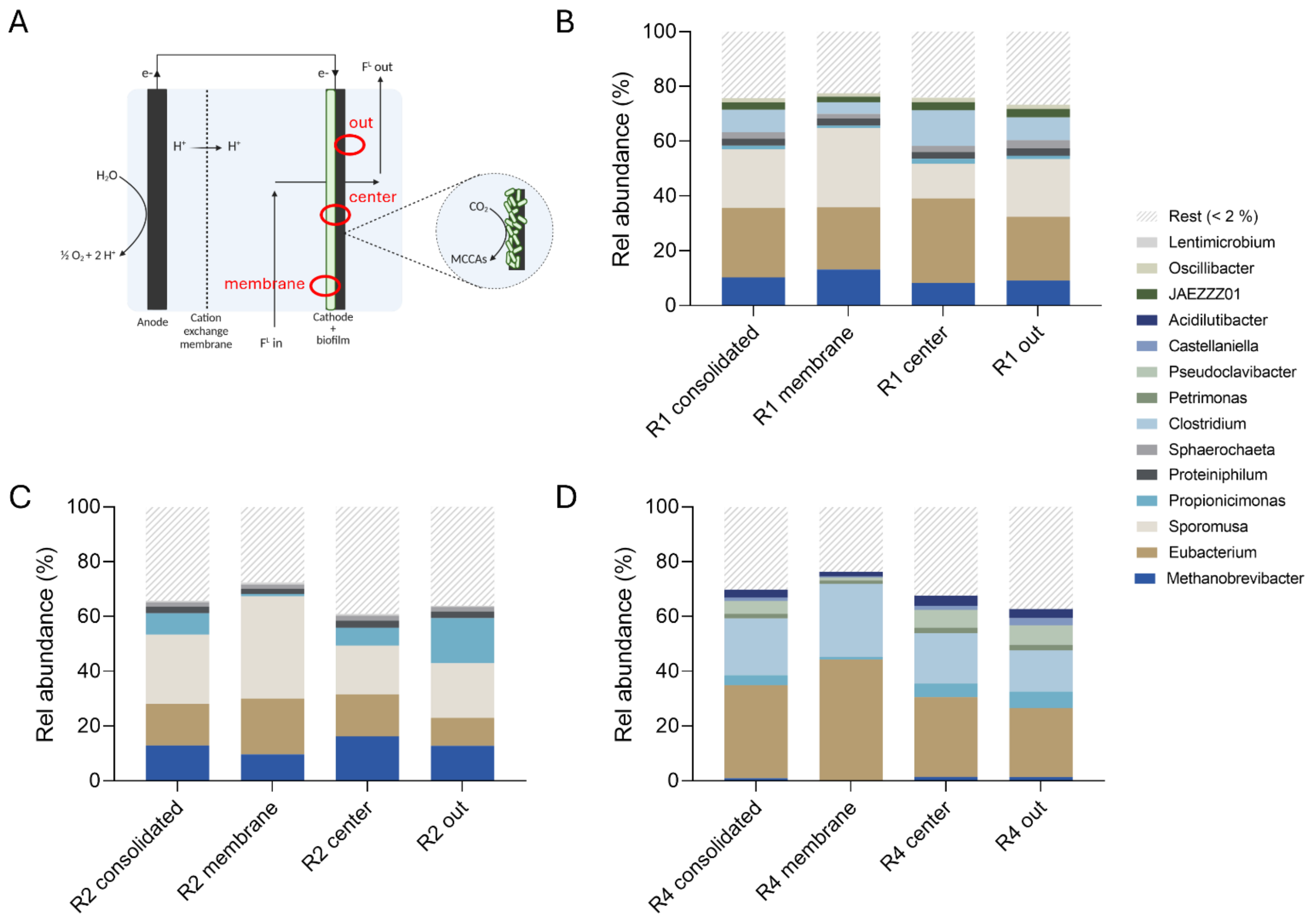


*Figure S2. Relative abundance of genera within the microbial communities of three MES reactors, R1 (B), R2 (C) and R4 (D) as according to metaproteomic data at three sampling locations (membrane, center and out) (A) and consolidated. The rest group includes the proteome assigned to bins that did not pass the checkM quality check (completeness > 90% and contamination <10%), bins that have a metagenomic abundance < 2% and the unbinned fraction.*

*Table S5. Total number of unique proteins in the metaproteome per reactor sample at the three locations (membrane, center, out) and consolidated.*

| Sample | membrane | center | out | consolidated |
|---|---|---|---|---|
| R1 | 4485 | 4179 | 4430 | **6756** |
| R2 | 3585 | 4306 | 4565 | **6789** |
| R4 | 3508 | 3698 | 2960 | **5968** |

*Table S6. The number of unique hypothetical proteins in the metaproteome per reactor sample at three locations (membrane, center, out) and consolidated.*

| Sample | membrane | center | out | consolidated |
|---|---|---|---|---|
| R1 | 1354 | 1176 | 1369 | **2001** |
| R2 | 1038 | 1255 | 1333 | **1924** |
| R4 | 891 | 932 | 821 | **1575** |

*Table S7. The number of uniquely identified proteins per MAG and the rest fraction, all three locations consolidated.*

| Number of unique identified proteins | | | |
|---|---|---|---|
| **MAG genus** | **R1** | **R2** | **R4** |
| Methanobrevibacter | 567 | 567 | 98 |
| Eubacterium | 811 | 568 | 989 |
| Sporomusa | 872 | 874 | |
| Propionicimonas | 162 | 756 | 297 |
| Proteiniphilum | 292 | 221 | |
| Sphaerochaeta | 143 | 97 | |
| Clostridium | 486 | | 736 |
| Petrimonas | | | 179 |
| Pseudoclavibacter | | | 246 |
| Pseudoclavibacter | | | 154 |
| Castellaniella | | | 118 |
| Acidilutibacter | | | 184 |
| JAEZZZ01 | 263 | | |
| Oscillibacter | 146 | | |
| Lentimicrobium | | 103 | |
| Rest (< 2 %) | 3014 | 3603 | 2967 |
| total | 6756 | 6789 | 5968 |

*Table S8. Overview of the manual search terms (KO IDs and EC numbers) that were used to functionally analyse the metagenomic and metaproteomic data. Open reading frames in the metagenomes were identified by Prokka and annotated by eggNOG-mapper V2. The search terms per pathway were retrieved from KEGG database [3].*

| Pathway | Description | KO ID / search term |
|---|---|---|
| Wood-Ljungdahl | EC 1.17.1.10: Formate dehydrogenase (NADP+) (FDH) (1) | 1.17.1.10 |
| Wood-Ljungdahl | EC 1.17.1.10: Formate dehydrogenase (NADP+) (FDH) (1) | K05299 |
| Wood-Ljungdahl | EC 1.17.1.10: Formate dehydrogenase (NADP+) (FDH) (1) | K15022 |
| Wood-Ljungdahl | EC 1.17.1.11: Formate dehydrogenase (FDH) bifurcating (NAD+, ferredoxin) (1) | 1.17.1.11 |
| Wood-Ljungdahl | EC 1.17.1.11: Formate dehydrogenase (FDH) bifurcating (NAD+, ferredoxin) (1) | K22338 |
| Wood-Ljungdahl | EC 1.17.1.11: Formate dehydrogenase (FDH) bifurcating (NAD+, ferredoxin) (1) | K22339 |
| Wood-Ljungdahl | EC 1.17.1.11: Formate dehydrogenase (FDH) bifurcating (NAD+, ferredoxin) (1) | K22340 |
| Wood-Ljungdahl | EC 1.17.1.11: Formate dehydrogenase (FDH) bifurcating (NAD+, ferredoxin) (1) | K22341 |
| Wood-Ljungdahl | EC 1.17.98.-: Formate dehydrogenase (FDH) (linked to hydrogenase 1.12.7.-) (1) | 1.17.98 |
| Wood-Ljungdahl | EC 1.17.98.-: Formate dehydrogenase (FDH) (linked to hydrogenase 1.12.7.-) (1) | 1.17.98.4 |
| Wood-Ljungdahl | EC 1.17.98.-: Formate dehydrogenase (FDH) (linked to hydrogenase 1.12.7.-) (1) | K22015 |
| Wood-Ljungdahl | EC 1.12.7.-: hydrogenase linked to 1.17.98.- (1) | K25123 |
| Wood-Ljungdahl | EC -: hydrogenase linked to 1.17.98.- (hycB) (1) | K25124 |
| Wood-Ljungdahl | EC 1.17.1.9: Formate dehydrogenase (NAD+) (FDH) (1) | 1.17.1.9 |
| Wood-Ljungdahl | EC 1.17.1.9: Formate dehydrogenase (NAD+) (FDH) (1) | K00122 |
| Wood-Ljungdahl | EC 1.17.1.9: Formate dehydrogenase (NAD+) (FDH) (1) | K00123 |
| Wood-Ljungdahl | EC 1.17.1.9: Formate dehydrogenase (NAD+) (FDH) (1) | K00126 |
| Wood-Ljungdahl | EC 1.17.1.9: Formate dehydrogenase (NAD+) (FDH) (1) | K22515 |
| Wood-Ljungdahl | EC 6.3.4.3: Formate-tetrahydrofolate ligase (FTS) (2) | 6.3.4.3 |
| Wood-Ljungdahl | EC 6.3.4.3: Formate-tetrahydrofolate ligase (FTS) (2) | K01938 |
| Wood-Ljungdahl | EC 3.5.4.9: methenyltetrahdyrofolate cyclohydrolase (MTC) (3) | 3.5.4.9 |
| Wood-Ljungdahl | EC 3.5.4.9: methenyltetrahdyrofolate cyclohydrolase (MTC) (3) | K01500 |
| Wood-Ljungdahl | EC 3.5.4.9/1.5.1.5: methenyltetrahdyrofolate cyclohydrolase / dehydrogenase (3 & 4) | K01491 |
| Wood-Ljungdahl | EC 3.5.4.9/1.5.1.15: methenyltetrahdyrofolate cyclohydrolase / dehydrogenase (3 & 4) | K13403 |
| Wood-Ljungdahl | EC 1.5.1.15: methylenetetrahydrofolate dehydrogenase (MTD) (4) | K00295 |
| Wood-Ljungdahl | EC 1.5.1.15: methylenetetrahydrofolate dehydrogenase (MTD) (4) | 1.5.1.15 |
| Wood-Ljungdahl | EC 1.5.1.5: methylenetetrahydrofolate dehydrogenase (MTD) (4) | 1.5.1.5 |
| Wood-Ljungdahl | EC 1.5.1.54: methylenetetrahydrofolate reductase (MTR) (NADH) (5) | 1.5.1.54 |
| Wood-Ljungdahl | EC 1.5.1.54: methylenetetrahydrofolate reductase (MTR) (NADH) (5) | K00297 |
| Wood-Ljungdahl | EC - :methylenetetrahydrofolate reductase (NADH) (metV) (MTR) (5) | K25007 |
| Wood-Ljungdahl | EC 7.-.-.-: methylenetetrahydrofolate reductase (NADH) (rnfC2) (MTR) (5) | K25008 |
| Wood-Ljungdahl | EC 1.5.7.1: methylenetetrahydrofolate reductase (MTR) (ferredoxin) (5) | 1.5.7.1 |
| Wood-Ljungdahl | EC 1.5.1.20: methylenetetrahydrofolate reductase (MTR) (NAD(P)H) (5) | 1.5.1.20 |
| Wood-Ljungdahl | EC 1.5.1.20: methylenetetrahydrofolate reductase (MTR) (NAD(P)H) (5) | K25009 |
| Wood-Ljungdahl | EC 2.1.1.258: methylTHF corrinoid/iron sulfur protein methyltransferase (MTF) (6) | 2.1.1.258 |
| Wood-Ljungdahl | EC 2.1.1.258: methylTHF corrinoid/iron sulfur protein methyltransferase (MTF) (6) | K15023 |
| Wood-Ljungdahl | EC 1.2.7.4: anaerobic carbon monoxide dehydrogenase (CODH) (7) | 1.2.7.4 |
| Wood-Ljungdahl | EC 1.2.7.4: anaerobic carbon monoxide dehydrogenase (CODH) (7) | K00198 |
| Wood-Ljungdahl | EC 1.2.7.4: anaerobic carbon monoxide dehydrogenase (CODH) (7) | K00196 |
| Wood-Ljungdahl | EC 2.3.1.169: CO-methylating acetyl-CoA synthase (ACS) (8) | 2.3.1.169 |
| Wood-Ljungdahl | EC 2.3.1.169: CO-methylating acetyl-CoA synthase (acsB) (ACS) (8) | K14138 |
| Wood-Ljungdahl | EC 2.1.1.245: CO-methylating acetyl-CoA synthase (ACS) (8) | 2.1.1.245 |
| Wood-Ljungdahl | EC 2.1.1.245: CO-methylating acetyl-CoA synthase (acsC) (ACS) (8) | K00197 |
| Wood-Ljungdahl | EC 2.1.1.245: CO-methylating acetyl-CoA synthase (acsD) (ACS) (8) | K00194 |
| Wood-Ljungdahl | EC 2.3.1.8: phosphate acetyltransferase (PTA) (9) | 2.3.1.8 |
| Wood-Ljungdahl | EC 2.3.1.8: phosphate acetyltransferase (PTA) (9) | K00625 |
| Wood-Ljungdahl | EC 2.3.1.8: phosphate acetyltransferase (PTA) (9) | K13788 |
| Wood-Ljungdahl | EC 2.3.1.8: phosphate acetyltransferase (PTA) (9) | K15024 |
| Wood-Ljungdahl | EC 2.7.2.1: acetate kinase (ACK) (10) | 2.7.2.1 |
| Wood-Ljungdahl | EC 2.7.2.1: acetate kinase (ACK) (10) | K00925 |
| Ethanol production | EC 1.2.7.5: aldehyde ferredoxin oxidoreductase (AOR) (11) | 1.2.7.5 |
| Ethanol production | EC 1.2.7.5: aldehyde ferredoxin oxidoreductase (AOR) (11) | k03738 |
| Ethanol production | EC 1.2.1.10: acetaldehyde dehydrogenase (ACDH) (12) | 1.2.1.10 |
| Ethanol production | EC 1.2.1.10: acetaldehyde dehydrogenase (ACDH) (12) | K00132 |
| Ethanol production | EC 1.2.1.10: acetaldehyde dehydrogenase (ACDH) (12) | K04073 |
| Ethanol production | EC 1.2.1.10 / 1.1.1.1: acetaldehyde dehydrogenase / alcohol dehydrogenase (ADH) (12 & 13) | K04072 |
| Ethanol production | EC 1.1.1.1: alcohol dehydrogenase (ADH) (13) | 1.1.1.1 |
| Ethanol production | EC 1.1.1.1: alcohol dehydrogenase (ADH) (13) | K00001 |
| Ethanol production | EC 1.1.1.1: alcohol dehydrogenase (ADH) (13) | K18857 |
| Ethanol production | EC 1.1.1.1: alcohol dehydrogenase (ADH) (13) | K13954 |
| Ethanol production | EC 1.1.1.1: alcohol dehydrogenase (ADH) (13) | K00121 |
| Ethanol production | EC 1.1.1.1: alcohol dehydrogenase (ADH) propanol-preferring (13) | K13953 |
| Ethanol production | EC 1.1.1.2: alcohol dehydrogenase (ADH) (NADP+) (13) | 1.1.1.2 |

| | | |
|---|---|---|
| Ethanol production | EC 1.1.1.2: alcohol dehydrogenase (ADH) (NADP+) (13) | K00002 |
| Lactate production | EC 1.2.7.1: pyruvate ferredoxin oxidoreductase (PFOR) (14) | 1.2.7.1 |
| Lactate production | EC 1.2.7.1: pyruvate ferredoxin oxidoreductase (PFOR) (14) | K00169 |
| Lactate production | EC 1.2.7.1: pyruvate ferredoxin oxidoreductase (PFOR) (14) | K00170 |
| Lactate production | EC 1.2.7.1: pyruvate ferredoxin oxidoreductase (PFOR) (14) | K00171 |
| Lactate production | EC 1.2.7.1: pyruvate ferredoxin oxidoreductase (PFOR) (14) | K00172 |
| Lactate production | EC 1.2.7.1: pyruvate ferredoxin oxidoreductase (PFOR) (14) | K03737 |
| Lactate production | EC 1.2.7.1: pyruvate ferredoxin oxidoreductase (PFOR) (14) | K00189 |
| Lactate production | EC 1.1.1.27: L-lactate dehydrogenase (LDH) (15) | 1.1.1.27 |
| Lactate production | EC 1.1.1.27: L-lactate dehydrogenase (LDH) (15) | K00016 |
| Lactate production | EC 1.1.1.436: lactate dehydrogenase electron bifurcating (LDH/Etf) (15) | 1.1.1.436 |
| Lactate production | EC 1.1.1.436: lactate dehydrogenase electron bifurcating (LDH/Etf) (15) | K26113 |
| Hydrogenotrophic methanogenesis | EC 1.2.7.12: formylmethanofuran dehydrogenase (1) | 1.2.7.12 |
| Hydrogenotrophic methanogenesis | EC 1.2.7.12: formylmethanofuran dehydrogenase (1) | K00200 |
| Hydrogenotrophic methanogenesis | EC 1.2.7.12: formylmethanofuran dehydrogenase (1) | K00201 |
| Hydrogenotrophic methanogenesis | EC 1.2.7.12: formylmethanofuran dehydrogenase (1) | K00202 |
| Hydrogenotrophic methanogenesis | EC 1.2.7.12: formylmethanofuran dehydrogenase (1) | K00203 |
| Hydrogenotrophic methanogenesis | EC 1.2.7.12: formylmethanofuran dehydrogenase (1) | K00204 |
| Hydrogenotrophic methanogenesis | EC 1.2.7.12: formylmethanofuran dehydrogenase (1) | K00205 |
| Hydrogenotrophic methanogenesis | EC 1.2.7.12: formylmethanofuran dehydrogenase (1) | K11261 |
| Hydrogenotrophic methanogenesis | EC 1.2.7.12: formylmethanofuran dehydrogenase (1) | K11260 |
| Hydrogenotrophic methanogenesis | EC 2.3.1.101: formylmethanofuran-tetrahydromethanopterin N-formyltransferase (2) | 2.3.1.101 |
| Hydrogenotrophic methanogenesis | EC 2.3.1.101: formylmethanofuran-tetrahydromethanopterin N-formyltransferase (2) | K00672 |
| Hydrogenotrophic methanogenesis | EC 3.5.4.27: methenyltetrahydromethanopterin cyclohydrolase (3) | 3.5.4.27 |
| Hydrogenotrophic methanogenesis | EC 3.5.4.27: methenyltetrahydromethanopterin cyclohydrolase (3) | K01499 |
| Hydrogenotrophic methanogenesis | EC 1.5.98.1: methylenetetrahydromethanopterin dehydrogenase F420 (4) | 1.5.98.1 |
| Hydrogenotrophic methanogenesis | EC 1.5.98.1: methylenetetrahydromethanopterin dehydrogenase F420 (4) | K00319 |
| Hydrogenotrophic methanogenesis | EC 1.12.98.2: methenyltetrahydromethanopterin dehydrogenase H2 (4) | 1.12.98.2 |
| Hydrogenotrophic methanogenesis | EC 1.12.98.2: methenyltetrahydromethanopterin dehydrogenase H2 (4) | K13942 |
| Hydrogenotrophic methanogenesis | EC 1.5.98.2: methylenetetrahydromethanopterin reductase (5) | 1.5.98.2 |
| Hydrogenotrophic methanogenesis | EC 1.5.98.2: methylenetetrahydromethanopterin reductase (5) | K00320 |
| Hydrogenotrophic methanogenesis | EC 7.2.1.4: tetrahydromethanopterin S-methyltransferase (6) | 7.2.1.4 |
| Hydrogenotrophic methanogenesis | EC 7.2.1.4: tetrahydromethanopterin S-methyltransferase (6) | K00577 |
| Hydrogenotrophic methanogenesis | EC 7.2.1.4: tetrahydromethanopterin S-methyltransferase (6) | K00578 |
| Hydrogenotrophic methanogenesis | EC 7.2.1.4: tetrahydromethanopterin S-methyltransferase (6) | K00579 |
| Hydrogenotrophic methanogenesis | EC 7.2.1.4: tetrahydromethanopterin S-methyltransferase (6) | K00580 |
| Hydrogenotrophic methanogenesis | EC 7.2.1.4: tetrahydromethanopterin S-methyltransferase (6) | K00581 |
| Hydrogenotrophic methanogenesis | EC 7.2.1.4: tetrahydromethanopterin S-methyltransferase (6) | K00582 |
| Hydrogenotrophic methanogenesis | EC 7.2.1.4: tetrahydromethanopterin S-methyltransferase (6) | K00583 |
| Hydrogenotrophic methanogenesis | EC 7.2.1.4: tetrahydromethanopterin S-methyltransferase (6) | K00584 |
| Hydrogenotrophic methanogenesis | EC 2.8.4.1: methyl-coenzyme M reductase (7) | 2.8.4.1 |
| Hydrogenotrophic methanogenesis | EC 2.8.4.1: methyl-coenzyme M reductase (7) | K00399 |
| Hydrogenotrophic methanogenesis | EC 2.8.4.1: methyl-coenzyme M reductase (7) | K00400 |
| Hydrogenotrophic methanogenesis | EC 2.8.4.1: methyl-coenzyme M reductase (7) | K00401 |
| Hydrogenotrophic methanogenesis | EC 2.8.4.1: methyl-coenzyme M reductase (7) | K00402 |
| Hydrogenotrophic methanogenesis | EC 2.8.4.1: methyl-coenzyme M reductase (7) | K03421 |
| Hydrogenotrophic methanogenesis | EC 2.8.4.1: methyl-coenzyme M reductase (7) | K03422 |
| Acetoclastic methanogenesis | EC 2.7.2.1: acetate kinase (1) | 2.7.2.1 |
| Acetoclastic methanogenesis | EC 2.7.2.1: acetate kinase (1) | K00925 |
| Acetoclastic methanogenesis | EC 2.3.1.8: phosphate acetyltransferase (2) | 2.3.1.8 |
| Acetoclastic methanogenesis | EC 2.3.1.8: phosphate acetyltransferase (2) | K00625 |
| Acetoclastic methanogenesis | EC 2.3.1.8: phosphate acetyltransferase (2) | K13788 |
| Acetoclastic methanogenesis | EC 6.2.1.1: acetyl-CoA synthetase (1 & 2) | 6.2.1.1 |
| Acetoclastic methanogenesis | EC 6.2.1.1: acetyl-CoA synthetase (1 & 2) | K01895 |
| Acetoclastic methanogenesis | EC 2.3.1.169: acetyl-CoA decarbonylase/synthetase (3) | 2.3.1.169 |
| Acetoclastic methanogenesis | EC 2.3.1.169: acetyl-CoA decarbonylase/synthetase (3) | K00193 |
| Acetoclastic methanogenesis | EC 2.1.1.245: acetyl-CoA decarbonylase/synthetase (3) | 2.1.1.245 |
| Acetoclastic methanogenesis | EC 2.1.1.245: acetyl-CoA decarbonylase/synthetase (3) | K00197 |
| Acetoclastic methanogenesis | EC 2.1.1.245: acetyl-CoA decarbonylase/synthetase (3) | K00194 |
| Acetoclastic methanogenesis | EC 7.2.1.4: tetrahydromethanopterin S-methyltransferase (4) | 2.1.1.86 |
| Acetoclastic methanogenesis | EC 7.2.1.4: tetrahydromethanopterin S-methyltransferase (4) | 7.2.1.4 |
| Acetoclastic methanogenesis | EC 7.2.1.4: tetrahydromethanopterin S-methyltransferase (4) | K00577 |
| Acetoclastic methanogenesis | EC 7.2.1.4: tetrahydromethanopterin S-methyltransferase (4) | K00578 |
| Acetoclastic methanogenesis | EC 7.2.1.4: tetrahydromethanopterin S-methyltransferase (4) | K00579 |
| Acetoclastic methanogenesis | EC 7.2.1.4: tetrahydromethanopterin S-methyltransferase (4) | K00580 |
| Acetoclastic methanogenesis | EC 7.2.1.4: tetrahydromethanopterin S-methyltransferase (4) | K00581 |
| Acetoclastic methanogenesis | EC 7.2.1.4: tetrahydromethanopterin S-methyltransferase (4) | K00582 |
| Acetoclastic methanogenesis | EC 7.2.1.4: tetrahydromethanopterin S-methyltransferase (4) | K00583 |
| Acetoclastic methanogenesis | EC 7.2.1.4: tetrahydromethanopterin S-methyltransferase (4) | K00584 |
| Acetoclastic methanogenesis | EC 2.8.4.1: methyl-coenzyme M reductase (5) | 2.8.4.1 |
| Acetoclastic methanogenesis | EC 2.8.4.1: methyl-coenzyme M reductase (5) | K00399 |

| | | |
|---|---|---|
| Acetoclastic methanogenesis | EC 2.8.4.1: methyl-coenzyme M reductase (5) | K00400 |
| Acetoclastic methanogenesis | EC 2.8.4.1: methyl-coenzyme M reductase (5) | K00401 |
| Acetoclastic methanogenesis | EC 2.8.4.1: methyl-coenzyme M reductase (5) | K00402 |
| Acetoclastic methanogenesis | EC 2.8.4.1: methyl-coenzyme M reductase (5) | K03421 |
| Acetoclastic methanogenesis | EC 2.8.4.1: methyl-coenzyme M reductase (5) | K03422 |
| Reverse beta-oxidation | EC 2.3.1.16: acetyl-Coa acyltransferase (THL) (1) | K00632 |
| Reverse beta-oxidation | EC 2.3.1.16: acetyl-Coa acyltransferase (THL) (1) | 2.3.1.16 |
| Reverse beta-oxidation | EC 2.3.1.16: acetyl-Coa acyltransferase (THL) (1) | K07508 |
| Reverse beta-oxidation | EC 2.3.1.16: acetyl-Coa acyltransferase (THL) (1) | K07509 |
| Reverse beta-oxidation | EC 2.3.1.16: acetyl-Coa acyltransferase (THL) (1) | K07513 |
| Reverse beta-oxidation | EC 2.3.1.9: acetyl-Coa acyltransferase (THL) (1) | 2.3.1.9 |
| Reverse beta-oxidation | EC 2.3.1.9: acetyl-Coa acyltransferase (THL) (1) | K00626 |
| Reverse beta-oxidation | EC 1.1.1.35: 3-hydroxyacyl-Coa dehydrogenase (HAD) (2) | K00022 |
| Reverse beta-oxidation | EC 1.1.1.35: 3-hydroxyacyl-Coa dehydrogenase (HAD) (2) | 1.1.1.35 |
| Reverse beta-oxidation | EC 1.1.1.35: 3-hydroxyacyl-Coa dehydrogenase (HAD) (2) | K07516 |
| Reverse beta-oxidation | EC 1.1.1.36: 3-hydroxyacyl-Coa dehydrogenase (HAD) (R) (2) | 1.1.1.36 |
| Reverse beta-oxidation | EC 1.1.1.36: 3-hydroxyacyl-Coa dehydrogenase (HAD) (R) (2) | K00023 |
| Reverse beta-oxidation | EC 1.1.1.157: 3-hydroxyacyl-Coa dehydrogenase (HAD) (2) | 1.1.1.157 |
| Reverse beta-oxidation | EC 1.1.1.157: 3-hydroxyacyl-Coa dehydrogenase (HAD) (2) | K00074 |
| Reverse beta-oxidation | EC 4.2.1.17: enoyl-Coa hydratase (CRT) (3) | 4.2.1.17 |
| Reverse beta-oxidation | EC 4.2.1.17: enoyl-Coa hydratase (CRT) (3) | K07511 |
| Reverse beta-oxidation | EC 4.2.1.17: enoyl-Coa hydratase (CRT) (3) | K13767 |
| Reverse beta-oxidation | EC 4.2.1.17: enoyl-Coa hydratase (CRT) (3) | K01692 |
| Reverse beta-oxidation | EC 4.2.1.17: enoyl-Coa hydratase (CRT) (3) | K07515 |
| Reverse beta-oxidation | EC 4.2.1.17: enoyl-Coa hydratase (CRT) (3) | K01715 |
| Reverse beta-oxidation | EC 4.2.1.150: enoyl-Coa hydratase (CRT) (3) | 4.2.1.150 |
| Reverse beta-oxidation | EC 4.2.1.55: enoyl-Coa hydratase (CRT) (R) (3) | 4.2.1.55 |
| Reverse beta-oxidation | EC 4.2.1.55: enoyl-Coa hydratase (CRT) (R) (3) | K17865 |
| Reverse beta-oxidation | EC 1.1.1.35 / 4.2.1.17: 3-hydroxyacyl-Coa dehydrogenase (HAD) / enoyl-Coa hydratase (CRT) (2&3) | K10527 |
| Reverse beta-oxidation | EC 1.1.1.35 / 4.2.1.17: 3-hydroxyacyl-Coa dehydrogenase (HAD) / enoyl-Coa hydratase (CRT) (2&3) | K01825 |
| Reverse beta-oxidation | EC 1.1.1.35 / 4.2.1.17: 3-hydroxyacyl-Coa dehydrogenase (HAD) / enoyl-Coa hydratase (CRT) (2&3) | K01782 |
| Reverse beta-oxidation | EC 1.1.1.35 / 4.2.1.17: 3-hydroxyacyl-Coa dehydrogenase (HAD) / enoyl-Coa hydratase (CRT) (2&3) | K07514 |
| Reverse beta-oxidation | EC 1.3.8.1: acyl-CoA dehydrogenase (short-chain) (ACD) (4) | 1.3.8.1 |
| Reverse beta-oxidation | EC 1.3.8.1: acyl- CoA dehydrogenase (short-chain) (ACD) (4) | K00248 |
| Reverse beta-oxidation | EC 1.3.8.7: acyl- CoA dehydrogenase (medium-chain) (ACD) (4) | K00248 |
| Reverse beta-oxidation | EC 1.3.8.7: acyl- CoA dehydrogenase (medium-chain) (ACD) (4) | 1.3.8.7 |
| Reverse beta-oxidation | EC 1.3.99.-: acyl- CoA dehydrogenase (ACD) (4) | K06445 |
| Reverse beta-oxidation | EC 1.3.1.109: acyl- CoA dehydrogenase (medium-chain) (ACD) (etf complex; NAD+, ferredoxin) (4) | 1.3.1.109 |
| Reverse beta-oxidation | EC 2.8.3.8: acetate- CoA transferase (ACAT) (5) | 2.8.3.8 |
| Reverse beta-oxidation | EC 2.8.3.8: acetate- CoA transferase (ACAT) (5) | 2.8.3.- |
| Reverse beta-oxidation | EC 2.8.3.8: acetate- CoA transferase (ACAT) (5) | K19709 |
| Reverse beta-oxidation | EC 2.8.3.8 / 2.8.3.9: acetate- CoA transferase (ACAT) (5) | K01034 |
| Reverse beta-oxidation | EC 2.8.3.8 / 2.8.3.9: acetate- CoA transferase (ACAT) (5) | K01035 |
| Reverse beta-oxidation | EC 2.8.3.9: acetate- CoA transferase (ACAT) (5) | 2.8.3.9 |
| Reverse beta-oxidation | EC 3.1.2.20: thioesterase (TES) (5) | 3.1.2.20 |
| Reverse beta-oxidation | EC 3.1.2.20: thioesterase (TES) (5) | K01073 |
| Reverse beta-oxidation | EC 2.3.1.19: phosphate butyryltransferase (PTB) (6) | 2.3.1.19 |
| Reverse beta-oxidation | EC 2.3.1.19: phosphate butyryltransferase (PTB) (6) | K00634 |
| Reverse beta-oxidation | EC 2.7.2.7: butyrate kinase (BUK) (7) | 2.7.2.7 |
| Reverse beta-oxidation | EC 2.7.2.7: butyrate kinase (BUK) (7) | K00929 |
| Reverse beta-oxidation | EC 2.7.2.18: fatty acid kinase (FAK) (7) | 2.7.2.18 |
| Reverse beta-oxidation | EC 2.7.2.18: fatty acid kinase (FAK) (7) | K07030 |
| Fatty acid biosynthesis | EC 2.3.1.39: s-malonyltransferase (initation) (fabD) | 2.3.1.39 |
| Fatty acid biosynthesis | EC 2.3.1.39: s-malonyltransferase (initation) (fabD) | K00645 |
| Fatty acid biosynthesis | EC 2.3.1.180: fabY / fabH (initation) | 2.3.1.180 |
| Fatty acid biosynthesis | EC 2.3.1.180: fabY / fabH (initation) | K18473 |
| Fatty acid biosynthesis | EC 2.3.1.180: fabY / fabH (initation) | K00648 |
| Fatty acid biosynthesis | fatty acid synthase (1 & 2 & 3 & 4 & 5) | K11533 |
| Fatty acid biosynthesis | EC 2.3.1.41: 3-oxoacyl-ACP synthase (FabB) (2) | K00647 |
| Fatty acid biosynthesis | EC 2.3.1.41: 3-oxoacyl-ACP synthase (FabB) (2) | 2.3.1.41 |
| Fatty acid biosynthesis | EC 2.3.1.179: 3-oxoacyl-ACP synthase (FabF) (2) | K09458 |
| Fatty acid biosynthesis | EC 2.3.1.179: 3-oxoacyl-ACP synthase (FabF) (2) | 2.3.1.179 |
| Fatty acid biosynthesis | EC 1.1.1.100: 3-oxoacyl-ACP reductase (FabG) (3) | 1.1.1.100 |
| Fatty acid biosynthesis | EC 1.1.1.100: 3-oxoacyl-ACP reductase (FabG) (3) | K00059 |
| Fatty acid biosynthesis | EC 4.2.1.59: 3-hydroxyacyl-ACP dehydratase (fabA/fabZ) (4) | K01716 |
| Fatty acid biosynthesis | EC 4.2.1.59: 3-hydroxyacyl-ACP dehydratase (fabA/fabZ) (4) | 4.2.1.59 |

| | | |
|---|---|---|
| Fatty acid biosynthesis | EC 4.2.1.59: 3-hydroxyacyl-ACP dehydratase (fabA/fabZ) (4) | K02372 |
| Fatty acid biosynthesis | EC 1.3.1.9: enoyl-ACP reductase (fabI/fabK/fabV) (5) | 1.3.1.9 |
| Fatty acid biosynthesis | EC 1.3.1.9 & 1.3.1.10: enoyl-ACP reductase (fabI) (5) | 1.3.1.10 |
| Fatty acid biosynthesis | EC 1.3.1.9 & 1.3.1.10: enoyl-ACP reductase (fabI) (5) | K00208 |
| Fatty acid biosynthesis | EC 1.3.1.9: enoyl-ACP reductase (fabI/fabK/fabV) (5) | K02371 |
| Fatty acid biosynthesis | EC 1.3.1.104: enoyl-ACP reductase (fabL) (5) | K10780 |
| Fatty acid biosynthesis | EC 1.3.1.9: enoyl-ACP reductase (fabI/fabK/fabV) (5) | K00209 |
| rTCA cycle | EC 2.3.3.8: ATP-citrate lyase (cycle 1) (1) | 2.3.3.8 |
| rTCA cycle | EC 2.3.3.8: ATP-citrate lyase (cycle 1) (1) | K01648 |
| rTCA cycle | EC 2.3.3.8: ATP-citrate lyase (cycle 1) (1) | K15230 |
| rTCA cycle | EC 2.3.3.8: ATP-citrate lyase (cycle 1) (1) | K15231 |
| rTCA cycle | EC 4.1.3.34: citryl-coa lyase (cycle 2) (1) | 4.1.3.34 |
| rTCA cycle | EC 4.1.3.34: citryl-coa lyase (cycle 2) (1) | K01644 |
| rTCA cycle | EC 4.1.3.34: citryl-coa lyase (cycle 2) (1) | K15234 |
| rTCA cycle | EC 1.1.1.37: malate dehydrogenase (2) | 1.1.1.37 |
| rTCA cycle | EC 1.1.1.37: malate dehydrogenase (2) | K00024 |
| rTCA cycle | EC 4.2.1.2: furmarase (3) | 4.2.1.2 |
| rTCA cycle | EC 4.2.1.2: furmarase (3) | K01675 |
| rTCA cycle | EC 4.2.1.2: furmarase (3) | K01676 |
| rTCA cycle | EC 4.2.1.2: furmarase (3) | K01677 |
| rTCA cycle | EC 4.2.1.2: furmarase (3) | K01678 |
| rTCA cycle | EC 4.2.1.2: furmarase (3) | K01679 |
| rTCA cycle | EC 4.2.1.2: furmarase (3) | K01774 |
| rTCA cycle | EC 1.3.1.6: furmarate reductase (NADH) (4) | 1.3.1.6 |
| rTCA cycle | EC 1.3.1.6: furmarate reductase (NADH) (4) | K18556 |
| rTCA cycle | EC 1.3.1.6: furmarate reductase (NADH) (4) | K18557 |
| rTCA cycle | EC 1.3.1.6: furmarate reductase (NADH) (4) | K18558 |
| rTCA cycle | EC 1.3.1.6: furmarate reductase (NADH) (4) | K18559 |
| rTCA cycle | EC 1.3.1.6: furmarate reductase (NADH) (4) | K18560 |
| rTCA cycle | EC 1.3.5.1: furmarate reductase (quinol) (4) | 1.3.5.1 |
| rTCA cycle | EC 1.3.5.1: furmarate reductase (quinol) (4) | K00239 |
| rTCA cycle | EC 1.3.5.1: furmarate reductase (quinol) (4) | K00240 |
| rTCA cycle | EC 1.3.5.1: furmarate reductase (quinol) (4) | K00241 |
| rTCA cycle | EC 1.3.5.1: furmarate reductase (quinol) (4) | K00242 |
| rTCA cycle | EC 1.3.5.1: furmarate reductase (quinol) (4) | K00244 |
| rTCA cycle | EC 1.3.5.1: furmarate reductase (quinol) (4) | K00245 |
| rTCA cycle | EC 1.3.5.1: furmarate reductase (quinol) (4) | K00246 |
| rTCA cycle | EC 1.3.5.1: furmarate reductase (quinol) (4) | K00247 |
| rTCA cycle | EC 6.2.1.5: succinyl-coa synthetase (5) | 6.2.1.5 |
| rTCA cycle | EC 6.2.1.5: succinyl-coa synthetase (5) | K01902 |
| rTCA cycle | EC 6.2.1.5: succinyl-coa synthetase (5) | K01903 |
| rTCA cycle | EC 1.2.7.3: 2-oxoglutarate syntase (6) | 1.2.7.3 |
| rTCA cycle | EC 1.2.7.3: 2-oxoglutarate syntase (6) | K00174 |
| rTCA cycle | EC 1.2.7.3: 2-oxoglutarate syntase (6) | K00175 |
| rTCA cycle | EC 1.2.7.3: 2-oxoglutarate syntase (6) | K00176 |
| rTCA cycle | EC 1.2.7.3: 2-oxoglutarate syntase (6) | K00177 |
| rTCA cycle | EC 1.1.1.42: isocitrate dehydrogenase (cycle 1) (7) | 1.1.1.42 |
| rTCA cycle | EC 1.1.1.42: isocitrate dehydrogenase (cycle 1) (7) | K00031 |
| rTCA cycle | EC 6.4.1.7: 2-oxoglutarate carboxylase (cycle 2) (7) | 6.4.1.7 |
| rTCA cycle | EC 6.4.1.7: 2-oxoglutarate carboxylase (cycle 2) (7) | K20140 |
| rTCA cycle | EC 6.4.1.7: 2-oxoglutarate carboxylase (cycle 2) (7) | K20141 |
| rTCA cycle | EC 1.1.1.42: oxalosuccinate reductase (cycle 2) (8) | 1.1.1.42 |
| rTCA cycle | EC 1.1.1.42: oxalosuccinate reductase (cycle 2) (8) | K00031 |
| rTCA cycle | EC 4.2.1.3: aconitate hydratase (cycle 1: 8 & 9; cycle 2: 9 & 10) | 4.2.1.3 |
| rTCA cycle | EC 4.2.1.3: aconitate hydratase (cycle 1: 8 & 9; cycle 2: 9 & 10) | K01681 |
| rTCA cycle | EC 4.2.1.3: aconitate hydratase (cycle 1: 8 & 9; cycle 2: 9 & 10) | K01682 |
| rTCA cycle | EC 4.2.1.3: aconitate hydratase (cycle 1: 8 & 9; cycle 2: 9 & 10) | K27802 |
| rTCA cycle | EC 6.2.1.18: citryl-coa synthetase (cycle 2: 11) | 6.2.1.18 |
| rTCA cycle | EC 6.2.1.18: citryl-coa synthetase (cycle 2: 11) | K15232 |
| rTCA cycle | EC 6.2.1.18: citryl-coa synthetase (cycle 2: 11) | K15233 |
| rTCA cycle | EC 6.4.1.1: pyruvate carboxylase (cycle 1: 11) | 6.4.1.1 |
| rTCA cycle | EC 6.4.1.1: pyruvate carboxylase (cycle 1: 11) | k01958 |
| rTCA cycle | EC 6.4.1.1: pyruvate carboxylase (cycle 1: 11) | k01959 |
| rTCA cycle | EC 6.4.1.1: pyruvate carboxylase (cycle 1: 11) | k01960 |
| rTCA cycle | EC 1.2.7.1: pyruvate ferredoxin oxidoreductase (cycle 1: 10; cycle 2: 12) | 1.2.7.1 |
| rTCA cycle | EC 1.2.7.1: pyruvate ferredoxin oxidoreductase (cycle 1: 10; cycle 2: 12) | K00169 |
| rTCA cycle | EC 1.2.7.1: pyruvate ferredoxin oxidoreductase (cycle 1: 10; cycle 2: 12) | K00170 |
| rTCA cycle | EC 1.2.7.1: pyruvate ferredoxin oxidoreductase (cycle 1: 10; cycle 2: 12) | K00171 |
| rTCA cycle | EC 1.2.7.1: pyruvate ferredoxin oxidoreductase (cycle 1: 10; cycle 2: 12) | K00172 |
| rTCA cycle | EC 1.2.7.1: pyruvate ferredoxin oxidoreductase (cycle 1: 10; cycle 2: 12) | K03737 |
| Calvin-Benson cycle | EC 2.7.1.19: phophoribulokinase (1) | 2.7.1.19 |

| | | |
|---|---|---|
| Calvin-Benson cycle | EC 2.7.1.19: phophoribulokinase (1) | K00855 |
| Calvin-Benson cycle | EC 4.1.1.39: RuBisCO (2) | 4.1.1.39 |
| Calvin-Benson cycle | EC 4.1.1.39: RuBisCO (2) | K01601 |
| Calvin-Benson cycle | EC 4.1.1.39: RuBisCO (2) | K01602 |
| Wood-Ljungdahl rGly | EC 2.1.2.10: glycine cleavage complex T protein (5) | 2.1.2.10 |
| Wood-Ljungdahl rGly | EC 2.1.2.10: glycine cleavage complex T protein (5) | K00605 |
| Wood-Ljungdahl rGly | EC 1.4.4.2: glycine cleavage complex P protein (5) | 1.4.4.2 |
| Wood-Ljungdahl rGly | EC 1.4.4.2: glycine cleavage complex P protein (5) | K00281 |
| Wood-Ljungdahl rGly | EC 1.4.4.2: glycine cleavage complex P protein (5) | K00282 |
| Wood-Ljungdahl rGly | EC 1.4.4.2: glycine cleavage complex P protein (5) | K00283 |
| Wood-Ljungdahl rGly | EC 1.8.1.4: glycine cleavage complex L protein (5) | 1.8.1.4 |
| Wood-Ljungdahl rGly | EC 1.8.1.4: glycine cleavage complex L protein (5) | K00382 |
| Wood-Ljungdahl rGly | EC 1.21.4.2: glycine reductase (thioredoxin) (6) | 1.21.4.2 |
| Wood-Ljungdahl rGly | EC 1.21.4.2: glycine reductase (thioredoxin) (6) | K10671 |
| Wood-Ljungdahl rGly | EC 1.21.4.2: glycine reductase (thioredoxin) (6) | K10672 |
| 3-Hydroxypropionate bicycle | EC 6.4.1.2: acetyl-coa carboxylase biotin carboxy carrier protein (1) | 6.4.1.2 |
| 3-Hydroxypropionate bicycle | EC 6.4.1.2: acetyl-coa carboxylase biotin carboxy carrier protein (1) | K02160 |
| 3-Hydroxypropionate bicycle | EC 6.3.4.14: acetyl-coa carboxylase biotin carboxylase subunit (1) | 6.3.4.14 |
| 3-Hydroxypropionate bicycle | EC 6.3.4.14: acetyl-coa carboxylase biotin carboxylase subunit (1) | K01961 |
| 3-Hydroxypropionate bicycle | EC 2.1.3.15: acetyl-coa carboxylase (1) | 2.1.3.15 |
| 3-Hydroxypropionate bicycle | EC 2.1.3.15: acetyl-coa carboxylase (1) | K01962 |
| 3-Hydroxypropionate bicycle | EC 2.1.3.15: acetyl-coa carboxylase (1) | K01963 |
| 3-Hydroxypropionate bicycle | EC 1.2.1.75: malonyl-coa reductase (NADP+) (2) | 1.2.1.75 |
| 3-Hydroxypropionate bicycle | EC 1.2.1.75: malonyl-coa reductase (NADP+) (2) | K15017 |
| 3-Hydroxypropionate bicycle | EC 1.1.1.298: 3-hydroxypropionate dehydrogenase (NADP+) (3) | 1.1.1.298 |
| 3-Hydroxypropionate bicycle | EC 1.1.1.298: 3-hydroxypropionate dehydrogenase (NADP+) (3) | K15039 |
| 3-Hydroxypropionate bicycle | EC 1.2.1.75/1.1.1.298: malonyl-coa reductase/3-hydroxypropionate dehydrogenase (NADP+) (2&3) | K14468 |
| 3-Hydroxypropionate bicycle | EC 6.2.1.36: 3-hydroxypropionyl-coa synthase (4) | 6.2.1.36 |
| 3-Hydroxypropionate bicycle | EC 6.2.1.36: 3-hydroxypropionyl-coa synthase (4) | K15018 |
| 3-Hydroxypropionate bicycle | EC 4.2.1.116: 3-hydroxypropionyl-coa dehydratase (5) | 4.2.1.116 |
| 3-Hydroxypropionate bicycle | EC 4.2.1.116: 3-hydroxypropionyl-coa dehydratase (5) | K15019 |
| 3-Hydroxypropionate bicycle | EC 1.3.1.84: acrylyl-coa reductase (NADPH) (6) | 1.3.1.84 |
| 3-Hydroxypropionate bicycle | EC 1.3.1.84: acrylyl-coa reductase (NADPH) (6) | K15020 |
| 3-Hydroxypropionate bicycle | EC 1.3.1.84/4.2.1.116/6.2.1.36: 3-hydroxypropionyl-coa synthase / dehydratase / acrylyl-coa reductase (4&5&6) | K14469 |
| 3-Hydroxypropionate bicycle | EC 6.4.1.3/2.3.1.15: propionyl-coa carboxylase (CO2 fixating step) (7) | K15052 |
| 3-Hydroxypropionate bicycle | EC 6.4.1.3/2.3.1.15: propionyl-coa carboxylase (CO2 fixating step) (7) | 6.4.1.3 |
| 3-Hydroxypropionate bicycle | EC 6.4.1.3/2.3.1.15: propionyl-coa carboxylase (CO2 fixating step) (7) | 2.1.3.15 |
| 3-Hydroxypropionate bicycle | EC 5.1.99.1: methyl malonyl-coa epimerase (8) | 5.1.99.1 |
| 3-Hydroxypropionate bicycle | EC 5.1.99.1: methyl malonyl-coa epimerase (8) | K05606 |
| 3-Hydroxypropionate bicycle | EC 5.4.99.2: methylmalonyl-coa mutase (9) | 5.4.99.2 |
| 3-Hydroxypropionate bicycle | EC 5.4.99.2: methylmalonyl-coa mutase (9) | K01848 |
| 3-Hydroxypropionate bicycle | EC 5.4.99.2: methylmalonyl-coa mutase (9) | K01849 |
| 3-Hydroxypropionate bicycle | EC 5.4.99.2: methylmalonyl-coa mutase (9) | K01847 |
| 3-Hydroxypropionate bicycle | EC 2.8.3.22: succinyl-coa:(s)-malate coa-transferase (10) | 2.8.3.22 |
| 3-Hydroxypropionate bicycle | EC 2.8.3.22: succinyl-coa:(s)-malate coa-transferase (10) | K14471 |
| 3-Hydroxypropionate bicycle | EC 2.8.3.22: succinyl-coa:(s)-malate coa-transferase (10) | K14472 |
| 3-Hydroxypropionate bicycle | EC 4.2.1.2: furmarate hydratase (10.2) | K01679 |
| 3-Hydroxypropionate bicycle | EC 4.2.1.2: furmarate hydratase (10.2) | 4.2.1.2 |
| 3-Hydroxypropionate bicycle | EC 1.3.5.1: succinate dehydrogenase flavoprotein (10.3) | 1.3.5.1 |
| 3-Hydroxypropionate bicycle | EC 1.3.5.1: succinate dehydrogenase flavoprotein (10.3) | K00239 |
| 3-Hydroxypropionate bicycle | EC 1.3.5.1: succinate dehydrogenase flavoprotein (10.3) | K00240 |
| 3-Hydroxypropionate bicycle | EC 1.3.5.1: succinate dehydrogenase flavoprotein (10.3) | K00241 |
| 3-Hydroxypropionate bicycle | EC 4.1.3.24: malyl-coa (11 & alt.11 & alt. 7) | 4.1.3.24 |
| 3-Hydroxypropionate bicycle | EC 4.1.3.25: (s) - cytramalyl-coa lyase (11 & alt.11 & alt. 7) | 4.1.3.25 |
| 3-Hydroxypropionate bicycle | EC 4.1.3.24/4.1.3.25: malyl-coa / (s) - cytramalyl-coa lyase (11 & alt.11 & alt. 7) | K08691 |
| 3-Hydroxypropionate bicycle | EC 4.2.1.148: 2-methylfumaryl-coa hydratase (alt. 8) | 4.2.1.148 |
| 3-Hydroxypropionate bicycle | EC 4.2.1.148: 2-methylfumaryl-coa hydratase (alt. 8) | K14449 |
| 3-Hydroxypropionate bicycle | EC 5.4.1.3: 2-methylfumaryl-coa isomerase (alt. 9) | 5.4.1.3 |
| 3-Hydroxypropionate bicycle | EC 5.4.1.3: 2-methylfumaryl-coa isomerase (alt. 9) | K14470 |
| 3-Hydroxypropionate bicycle | EC 4.2.1.153: 3-methylfumaryl-coa hydratase (alt.10) | 4.2.1.153 |
| 3-Hydroxypropionate bicycle | EC 4.2.1.153: 3-methylfumaryl-coa hydratase (alt.10) | K09709 |
| 3-HP / 4-HB cycle | EC 6.4.1.2: acetyl-coa carboxylase (1) | 6.4.1.2 |
| 3-HP / 4-HB cycle | EC 6.4.1.2: acetyl-coa carboxylase (1) | K01962 |
| 3-HP / 4-HB cycle | EC 6.4.1.2: acetyl-coa carboxylase (1) | K01961 |
| 3-HP / 4-HB cycle | EC 6.4.1.2: acetyl-coa carboxylase (1) | K01963 |
| 3-HP / 4-HB cycle | EC 6.4.1.2: acetyl-coa carboxylase (1) | K18603 |
| 3-HP / 4-HB cycle | EC 6.4.1.2: acetyl-coa carboxylase (1) | K15036 |
| 3-HP / 4-HB cycle | EC 6.4.1.2: acetyl-coa carboxylase (1) | K18604 |
| 3-HP / 4-HB cycle | EC 1.2.1.75: malonyl-coa reductase (2) | 1.2.1.75 |

| | | |
|---|---|---|
| 3-HP / 4-HB cycle | EC 1.2.1.75: malonyl-coa reductase (2) | K15017 |
| 3-HP / 4-HB cycle | EC 1.2.1.75 / 1.1.1.298: malonyl-coa reductase / 3-hydroxypropionate dehydrogenase (3HP-4HB) (2 & 3) | K14468 |
| 3-HP / 4-HB cycle | EC 1.1.1.298: 3-hydroxypropionate dehydrogenase (3HP-4HB) (NADP+) (3) | 1.1.1.298 |
| 3-HP / 4-HB cycle | EC 1.1.1.298: 3-hydroxypropionate dehydrogenase (3HP-4HB) (NADP+) (3) | K15039 |
| 3-HP / 4-HB cycle | EC 6.2.1.36: 3-hydroxypropionyl-coenzyme A synthetase (4) | 6.2.1.36 |
| 3-HP / 4-HB cycle | EC 6.2.1.36: 3-hydroxypropionyl-coenzyme A synthetase (4) | K15018 |
| 3-HP / 4-HB cycle | EC 4.2.1.116: 3-hydroxypropionyl-coenzyme A dehydratase (5) | 4.2.1.116 |
| 3-HP / 4-HB cycle | EC 4.2.1.116: 3-hydroxypropionyl-coenzyme A dehydratase (5) | K15019 |
| 3-HP / 4-HB cycle | EC 1.3.1.84: acryloyl-coenzyme A reductase (6) | 1.3.1.84 |
| 3-HP / 4-HB cycle | EC 1.3.1.84: acryloyl-coenzyme A reductase (6) | K15020 |
| 3-HP / 4-HB cycle | EC 6.4.1.3: propionyl-coa carboxylase (7) | 6.4.1.3 |
| 3-HP / 4-HB cycle | EC -: propionyl-coa carboxylase (biotin carboxyl carrier protein) (7) | K15037 |
| 3-HP / 4-HB cycle | EC 6.4.1.2/6.4.1.3: acetyl-coa/propionyl-coa carboxylase (2 & 7) | K01964 |
| 3-HP / 4-HB cycle | EC 6.4.1.2/6.4.1.3: acetyl-coa/propionyl-coa carboxylase (2 & 7) | K15036 |
| 3-HP / 4-HB cycle | EC 5.1.99.1: methylmalonyl-coa epimerase (8) | 5.1.99.1 |
| 3-HP / 4-HB cycle | EC 5.1.99.1: methylmalonyl-coa epimerase (8) | K05606 |
| 3-HP / 4-HB cycle | EC 5.4.99.2: methylmalonyl-coa mutase (3HP-4HB) (9) | 5.4.99.2 |
| 3-HP / 4-HB cycle | EC 5.4.99.2: methylmalonyl-coa mutase (3HP-4HB) (9) | K01848 |
| 3-HP / 4-HB cycle | EC 5.4.99.2: methylmalonyl-coa mutase (3HP-4HB) (9) | K01849 |
| 3-HP / 4-HB cycle | EC 1.2.1.76: succinyl-coa reductase (10) | 1.2.1.76 |
| 3-HP / 4-HB cycle | EC 1.2.1.76: succinyl-coa reductase (10) | K15038 |
| 3-HP / 4-HB cycle | EC 1.2.1.76: succinyl-coa reductase (10) | K15017 |
| 3-HP / 4-HB cycle | EC 1.1.1.-: succinate semialdehyde reductase (NADPH) (11) | 1.1.1.- |
| 3-HP / 4-HB cycle | EC 1.1.1.-: succinate semialdehyde reductase (NADPH) (11) | K14465 |
| 3-HP / 4-HB cycle | EC 6.2.1.40: 4-hydroxybutyrate--coa ligase (AMP-forming) (12) | 6.2.1.40 |
| 3-HP / 4-HB cycle | EC 6.2.1.40: 4-hydroxybutyrate--coa ligase (AMP-forming) (12) | K14466 |
| 3-HP / 4-HB cycle | EC 6.2.1.40: 4-hydroxybutyrate--coa ligase (AMP-forming) (12) | K18861 |
| 3-HP / 4-HB cycle | EC 4.2.1.120: 4-hydroxybutytyl-coa dehydratase (13) | 4.2.1.120 |
| 3-HP / 4-HB cycle | EC 4.2.1.120: 4-hydroxybutytyl-coa dehydratase (13) | K14534 |
| 3-HP / 4-HB cycle | EC 4.2.1.17: 3-hydroxybutytyl-coa dehydratase (14) | 4.2.1.17 |
| 3-HP / 4-HB cycle | EC 1.1.1.35: 3-hydroxyacyl-coa dehydrogenase (15) | 1.1.1.35 |
| 3-HP / 4-HB cycle | EC 4.2.1.17/1.1.1.35: 3-hydroxybutyryl-CoA dehydratase / 3-hydroxyacyl-CoA dehydrogenase (14 & 15) | K15016 |
| 3-HP / 4-HB cycle | EC 2.3.1.9: acetyl-coa c-acetyltransferase (16) | 2.3.1.9 |
| 3-HP / 4-HB cycle | EC 2.3.1.9: acetyl-coa c-acetyltransferase (16) | K00626 |
| DC / 4-HB cycle | EC 1.2.7.1: pyruvate ferredoxin oxidoreductase (1) | 1.2.7.1 |
| DC / 4-HB cycle | EC 1.2.7.1: pyruvate ferredoxin oxidoreductase (1) | K00169 |
| DC / 4-HB cycle | EC 1.2.7.1: pyruvate ferredoxin oxidoreductase (1) | K00170 |
| DC / 4-HB cycle | EC 1.2.7.1: pyruvate ferredoxin oxidoreductase (1) | K00171 |
| DC / 4-HB cycle | EC 1.2.7.1: pyruvate ferredoxin oxidoreductase (1) | K00172 |
| DC / 4-HB cycle | EC 2.7.9.2: pyruvate, water dikinase (2) | 2.7.9.2 |
| DC / 4-HB cycle | EC 2.7.9.2: pyruvate, water dikinase (2) | K01007 |
| DC / 4-HB cycle | EC 4.1.1.31: phosphoenolpyruvate carboxylase (3) | 4.1.1.31 |
| DC / 4-HB cycle | EC 4.1.1.31: phosphoenolpyruvate carboxylase (3) | K01595 |
| DC / 4-HB cycle | EC 1.1.1.37: malate dehydrogenase (4) | 1.1.1.37 |
| DC / 4-HB cycle | EC 1.1.1.37: malate dehydrogenase (4) | K00024 |
| DC / 4-HB cycle | EC 4.2.1.2: fumarate hydratase (5) | 4.2.1.2 |
| DC / 4-HB cycle | EC 4.2.1.2: fumarate hydratase (5) | K01676 |
| DC / 4-HB cycle | EC 4.2.1.2: fumarate hydratase (5) | K01677 |
| DC / 4-HB cycle | EC 4.2.1.2: fumarate hydratase (5) | K01678 |
| DC / 4-HB cycle | EC 1.3.5.1: succinate dehydrogenase (6) | 1.3.5.1 |
| DC / 4-HB cycle | EC 1.3.5.1: succinate dehydrogenase (6) | K00239 |
| DC / 4-HB cycle | EC 1.3.5.1: succinate dehydrogenase (6) | K00240 |
| DC / 4-HB cycle | EC 1.3.5.1: succinate dehydrogenase (6) | K00241 |
| DC / 4-HB cycle | EC 1.3.5.1: succinate dehydrogenase (6) | K18860 |
| DC / 4-HB cycle | EC 6.2.1.5: succinyl-coa synthetase (7) | 6.2.1.5 |
| DC / 4-HB cycle | EC 6.2.1.5: succinyl-coa synthetase (7) | K01902 |
| DC / 4-HB cycle | EC 6.2.1.5: succinyl-coa synthetase (7) | K01903 |
| DC / 4-HB cycle | EC 1.2.1.76: succinyl-coa reductase (8) | 1.2.1.76 |
| DC / 4-HB cycle | EC 1.2.1.76: succinyl-coa reductase (8) | K15017 |
| DC / 4-HB cycle | EC 1.2.1.76: succinyl-coa reductase (8) | K15038 |
| DC / 4-HB cycle | EC 1.1.1.-: succinate semialdehyde reductase (NADPH) (9) | 1.1.1.- |
| DC / 4-HB cycle | EC 1.1.1.-: succinate semialdehyde reductase (NADPH) (9) | K14465 |
| DC / 4-HB cycle | EC 6.2.1.40: 4-hydroxybutyrate--coa ligase (AMP-forming) (10) | 6.2.1.40 |
| DC / 4-HB cycle | EC 6.2.1.40: 4-hydroxybutyrate--coa ligase (AMP-forming) (10) | K14467 |
| DC / 4-HB cycle | EC 6.2.1.40: 4-hydroxybutyrate--coa ligase (AMP-forming) (10) | K18861 |
| DC / 4-HB cycle | EC 4.2.1.120: 4-hydroxybutyryl-coa dehydratase (11) | 4.2.1.120 |
| DC / 4-HB cycle | EC 4.2.1.120: 4-hydroxybutyryl-coa dehydratase (11) | K14534 |
| DC / 4-HB cycle | EC 4.2.1.17: 3-hydroxygutytyl-coa dehydratase (12) | 4.2.1.17 |
| DC / 4-HB cycle | EC 1.1.1.35: 3-hydroxyacyl-coa dehydrogenase (13) | 1.1.1.35 |

| | | |
|---|---|---|
| DC / 4-HB cycle | EC 4.2.1.17/1.1.1.35: 3-hydroxygutytyl-coa dehydratase / 3-hydroxyacyl-coa dehydrogenase (12 & 13) | K15016 |
| DC / 4-HB cycle | EC 2.3.1.9: acetyl-CoA C-acetyltransferase (14) | 2.3.1.9 |
| DC / 4-HB cycle | EC 2.3.1.9: acetyl-CoA C-acetyltransferase (14) | K00626 |

#### 5. Functional proteomics abundance at different location samples

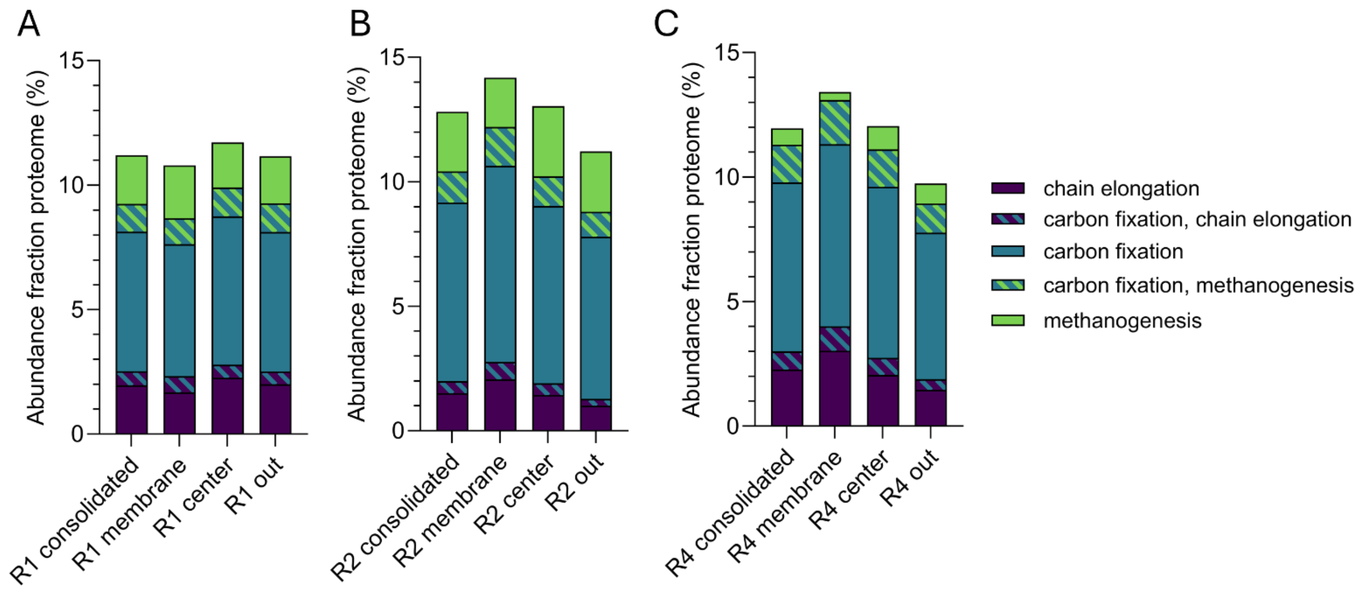


*Figure S3. Relative proteome abundance of carbon fixation enzymes to acetate, ethanol or lactate production (light blue), chain elongation from acetate to butyrate and caproate (dark blue) and methanogenesis (green) in reactors R1 (A), R2 (B) and R4 (C) at the different proteome sampling locations (membrane, center and out). A manual search was performed to functionally annotate the proteome to the three categories and can be found in supplementary Table S8.*

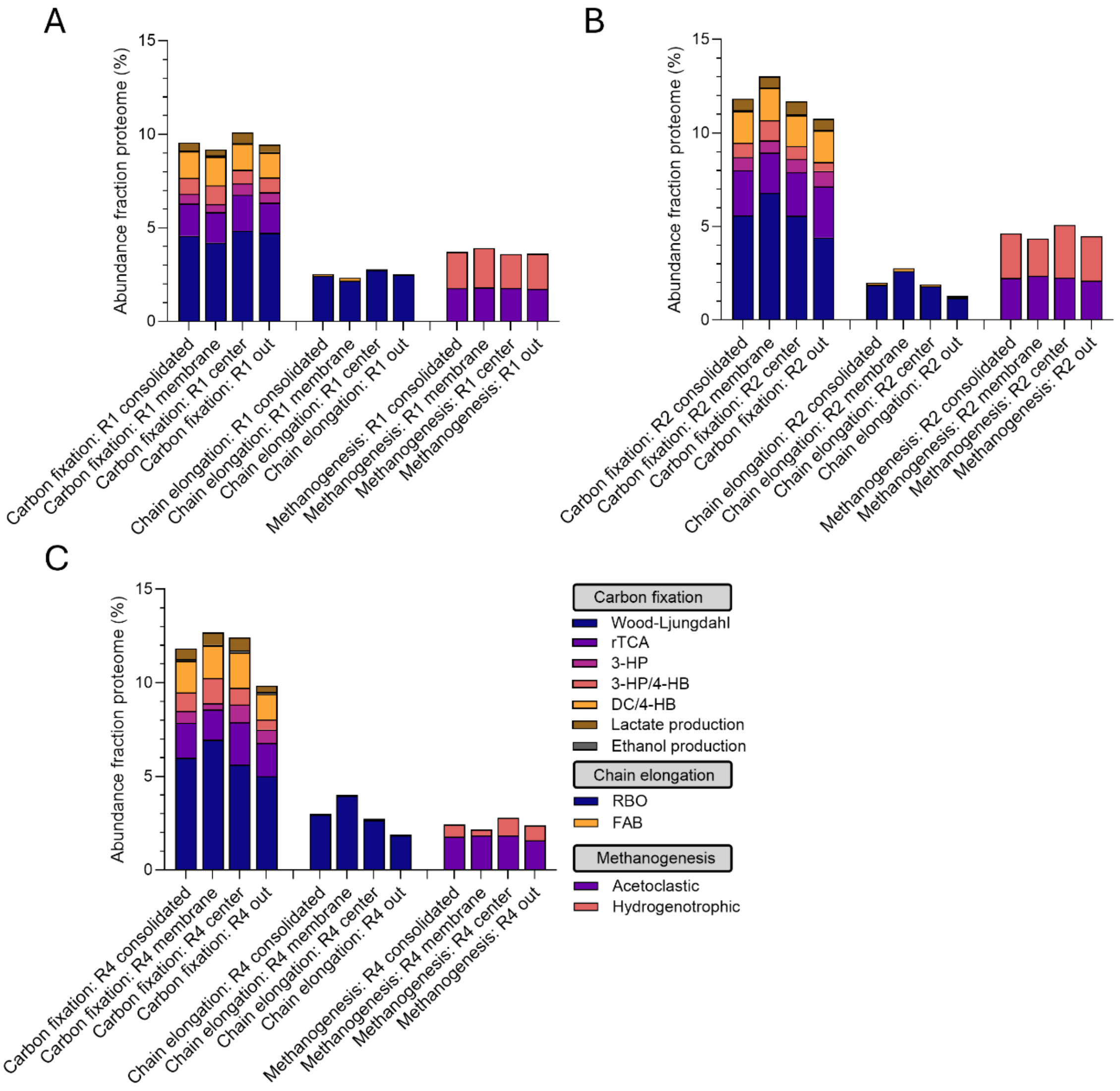


*Figure S4. Relative abundance of proteins in metaproteome of three categories (I) different carbon fixation routes to acetate, ethanol (from acetyl-CoA) or lactate (from acetyl-CoA), (II) chain elongation and (III) methanogenesis in reactors R1 (A), R2 (B) and R4 (C) at the different proteome sample locations (membrane, center and out). Overlapping enzymes between different pathways are accounted for in the fraction of each pathway, therefore the sum of the abundance of all pathways is slightly higher than the total abundance fraction of that category. A manual search was performed to functionally annotate the proteome to the three categories and can be found in supplementary Table S8.*

### 6. Protein expression WLP at membrane, center and out locations

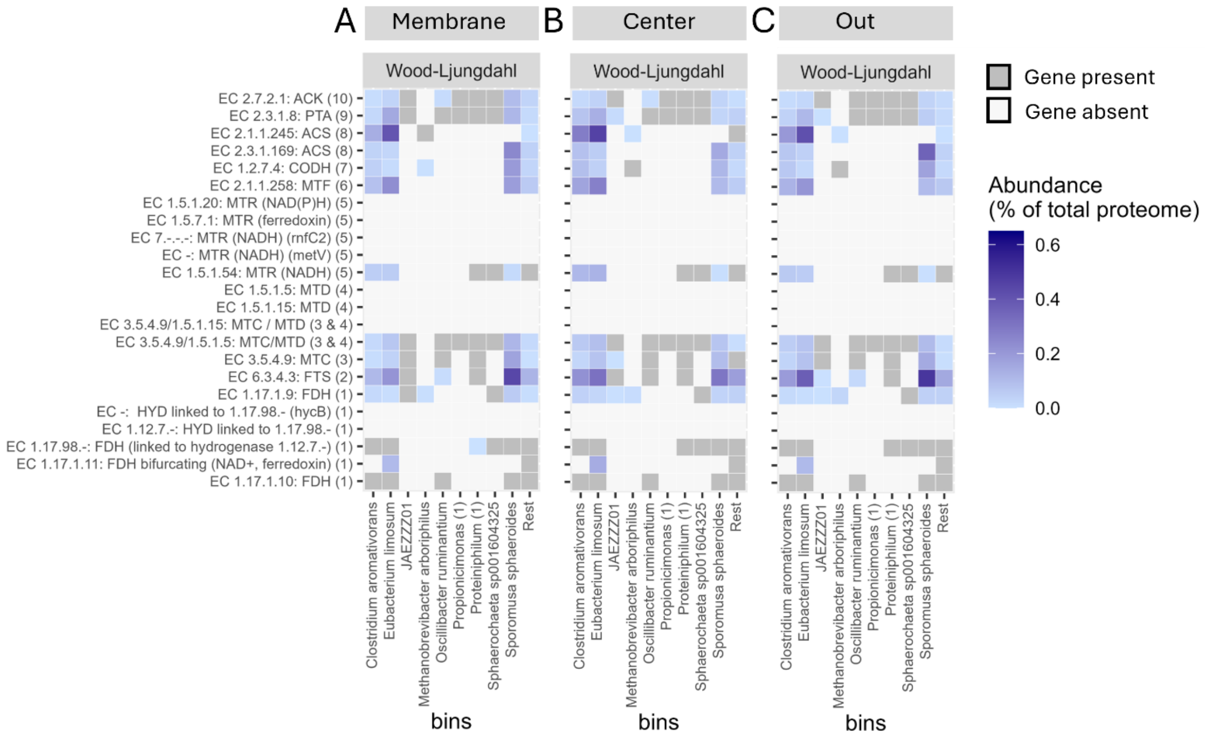


*Figure S5. Heatmaps showing the gene presence and protein abundance of the individual steps of the Wood-Ljungdahl Pathway (WLP) per MAG at the membrane (A), center (B) and out (C) proteome sampling locations in R1. Gene presence is shown in dark grey and protein abundance (%) in light to dark blue gradient.*

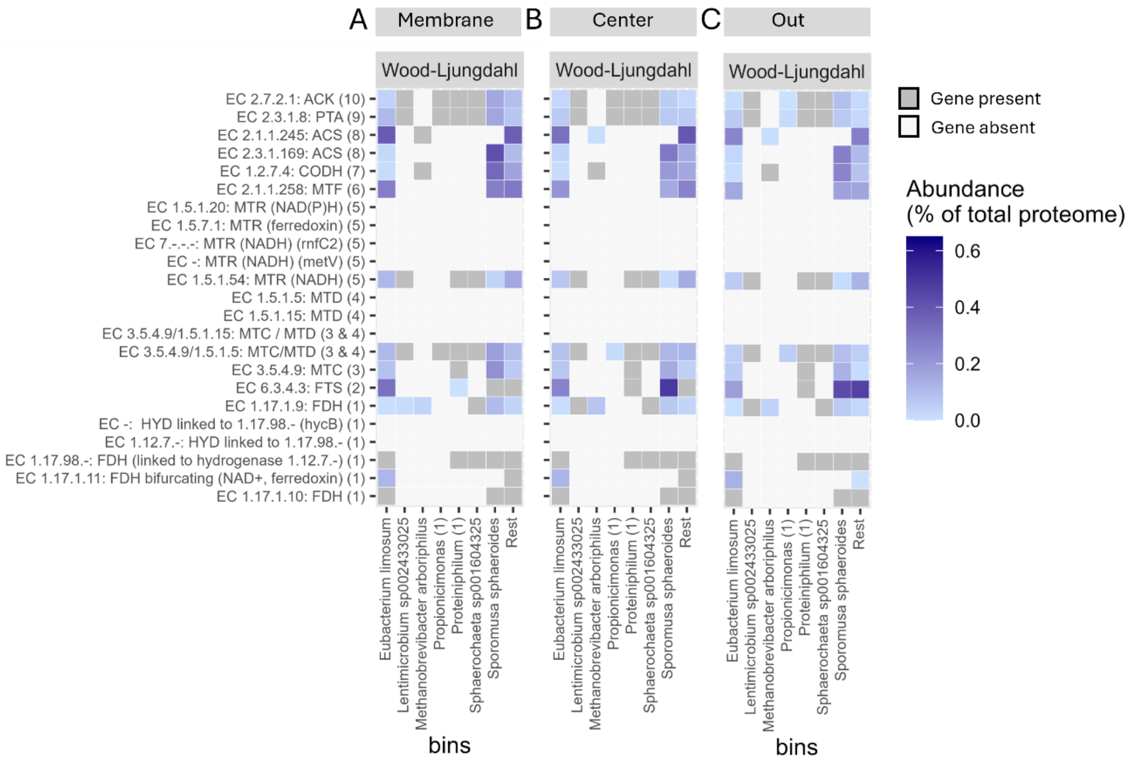


*Figure S6. Heatmaps showing the gene presence and protein abundance of the individual steps of the Wood-Ljungdahl Pathway (WLP) per MAG at the membrane (A), center (B) and out (C) proteome sampling locations in reactor 2 (R2). Gene presence is shown in dark grey and protein abundance (%) in light to dark blue gradient.*

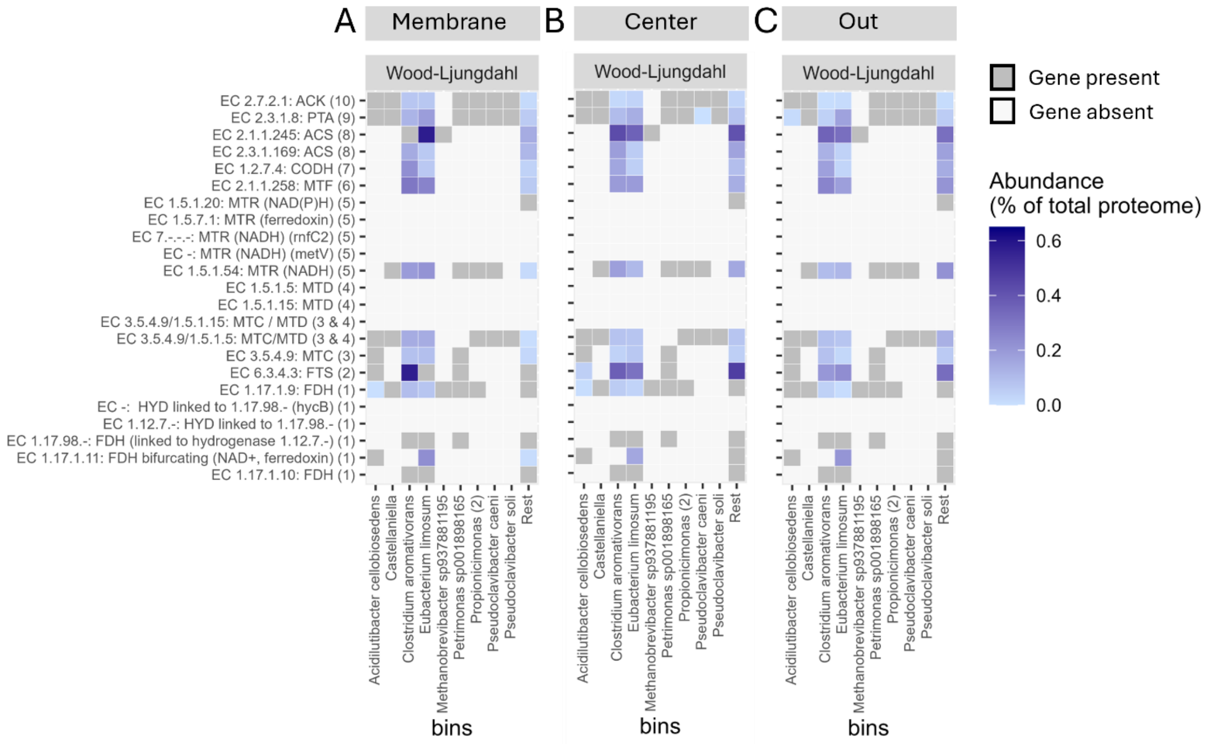


*Figure S7. Heatmaps showing the gene presence and protein abundance of the individual steps of the Wood-Ljungdahl Pathway (WLP) per MAG at the membrane (A), center (B) and out (C) proteome sampling locations in (reactor 4) R4. Gene presence is shown in dark grey and protein abundance (%) in light to dark blue gradient.*

#### 7. Protein expression ethanol and lactate production at membrane, center and out locations

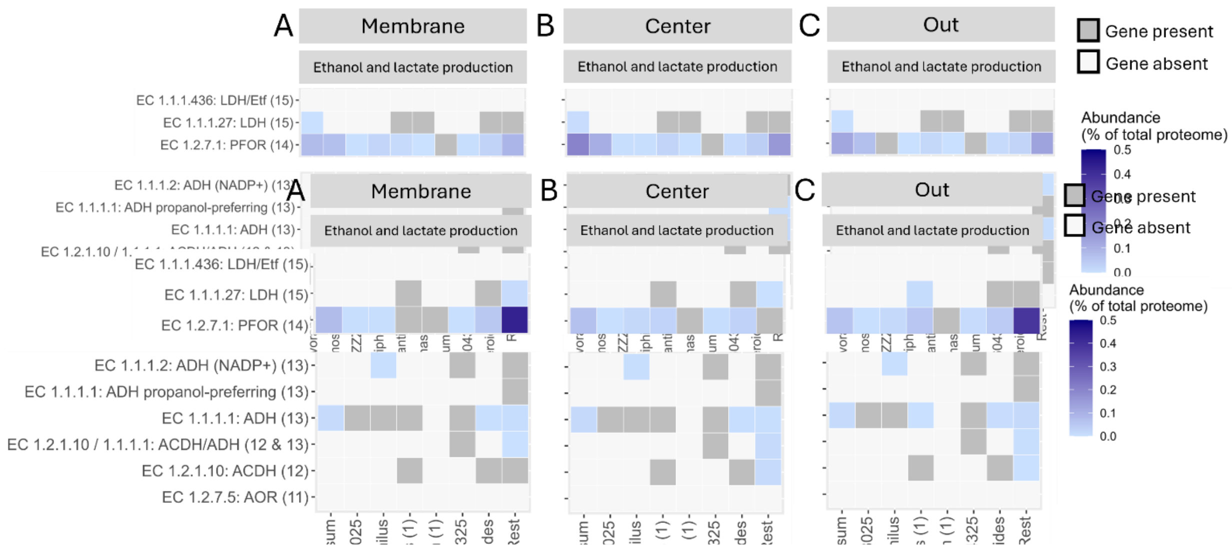


*Figure S8. Heatmaps showing the gene presence and protein abundance of the individual steps of ethanol and lactate production from acetyl-CoA per MAG at the membrane (A), center (B) and out (C) proteome sampling locations in reactor 1 (R1). Gene presence is shown in dark grey and protein abundance (%) in light to dark blue gradient.*

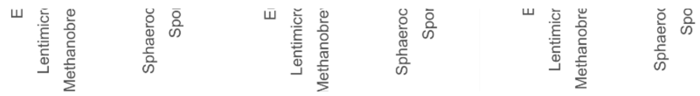

*Figure S9. Heatmaps showing the gene presence and protein abundance of the individual steps of ethanol and lactate production from acetyl-CoA per MAG at the membrane (A), center (B) and out (C) proteome sampling locations in reactor 2 (R2). Gene presence is shown in dark grey and protein abundance (%) in light to dark blue gradient.*

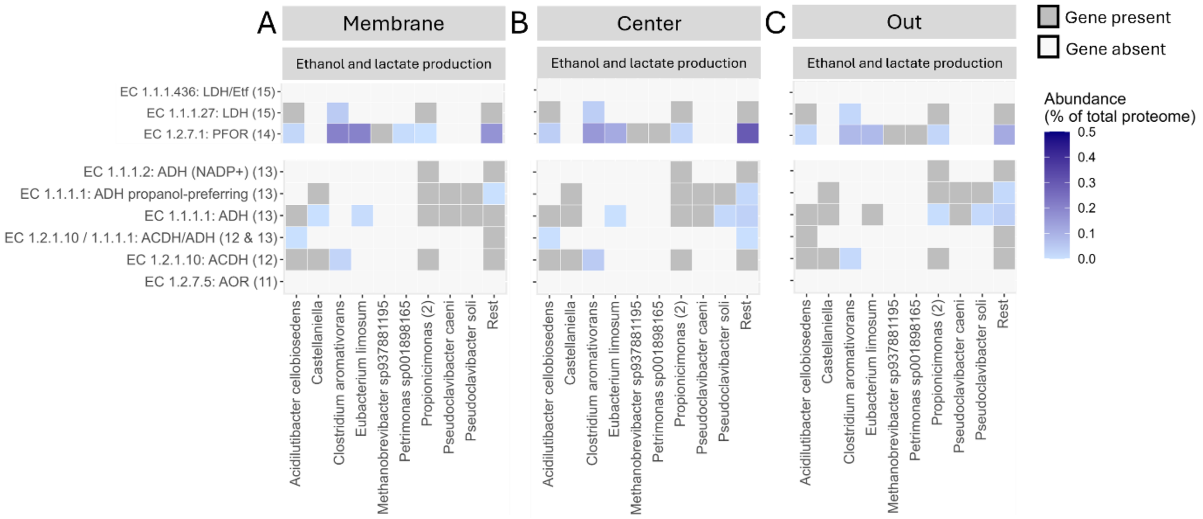


*Figure S10. Heatmaps showing the gene presence and protein abundance of the individual steps of ethanol and lactate production from acetyl-CoA per MAG at the membrane (A), center (B) and out (C) proteome sampling locations in reactor 4 (R4). Gene presence is shown in dark grey and protein abundance (%) in light to dark blue gradient.*

#### 8. Protein expression RBO at membrane, center and out locations

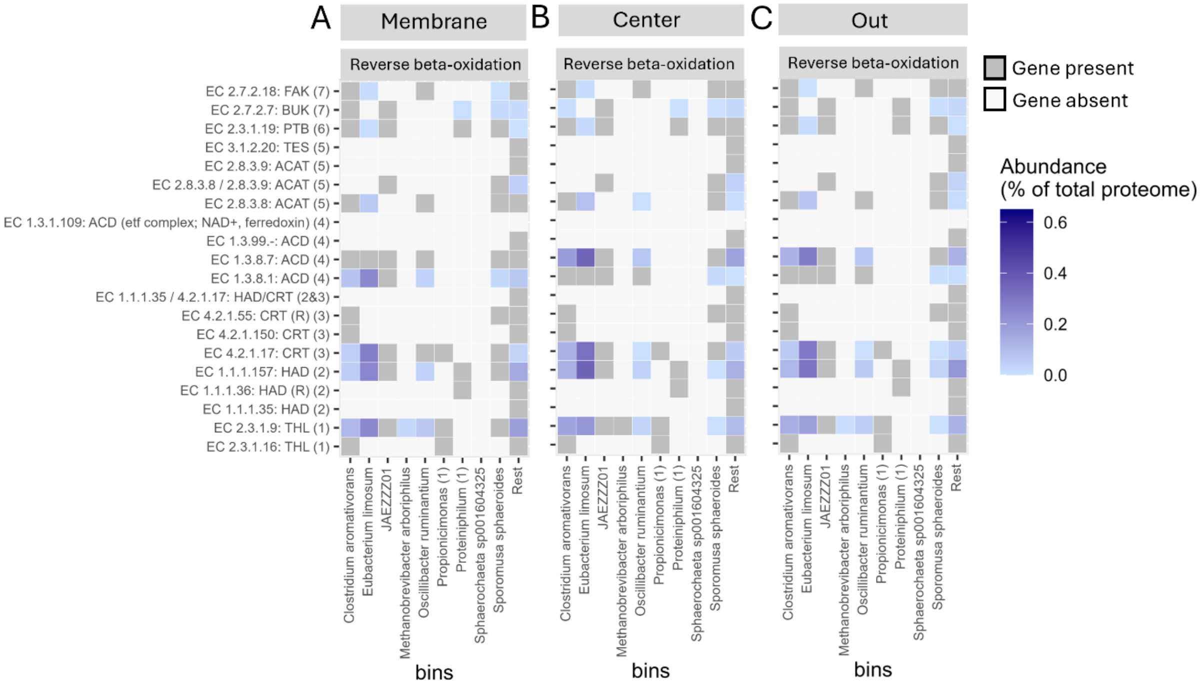


*Figure S11. Heatmaps showing the gene presence and protein abundance of the individual steps of the Reverse Beta-oxidation (RBO) per MAG at the membrane (A), center (B) and out (C) proteome sampling locations in reactor 1 (R1). Gene presence is shown in dark grey and protein abundance (%) in light to dark blue gradient.*

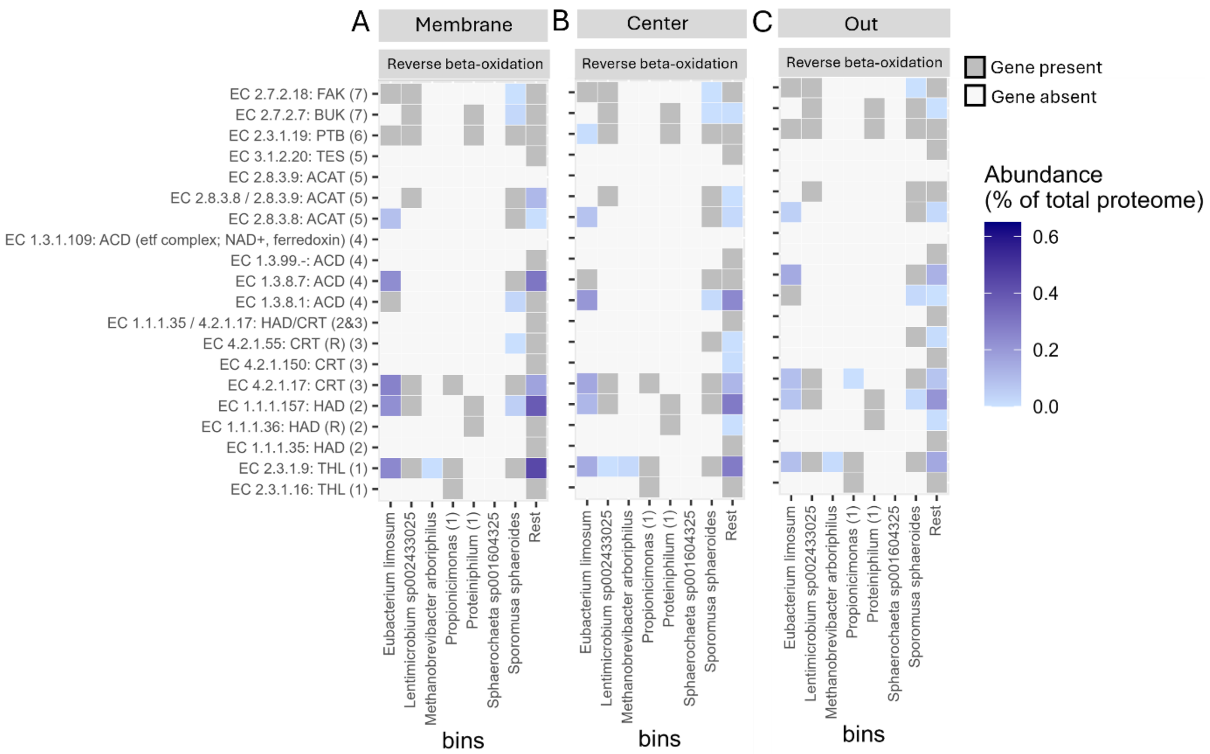


*Figure S12. Heatmaps showing the gene presence and protein abundance of the individual steps of the Reverse Beta-Oxidation (RBO) per MAG at the membrane (A), center (B) and out (C) proteome sampling locations in reactor 2 (R2). Gene presence is shown in dark grey and protein abundance (%) in light to dark blue gradient.*

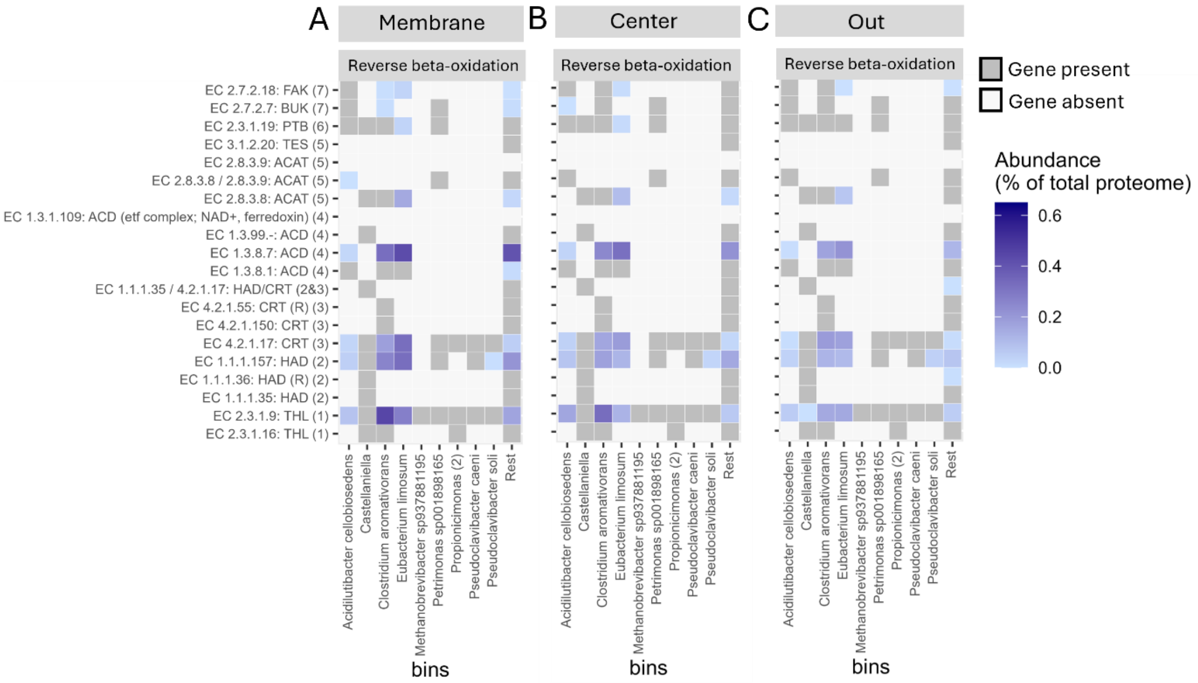


*Figure S13. Heatmaps showing the gene presence and protein abundance of the individual steps of the Reverse beta-oxidation (RBO) per MAG at the membrane (A), center (B) and out (C) proteome sampling locations in reactor 4 (R4). Gene presence is shown in dark grey and protein abundance (%) in light to dark blue gradient.*

#### 9. Protein expression of alternative carbon fixation routes, chain elongation and methanogenesis

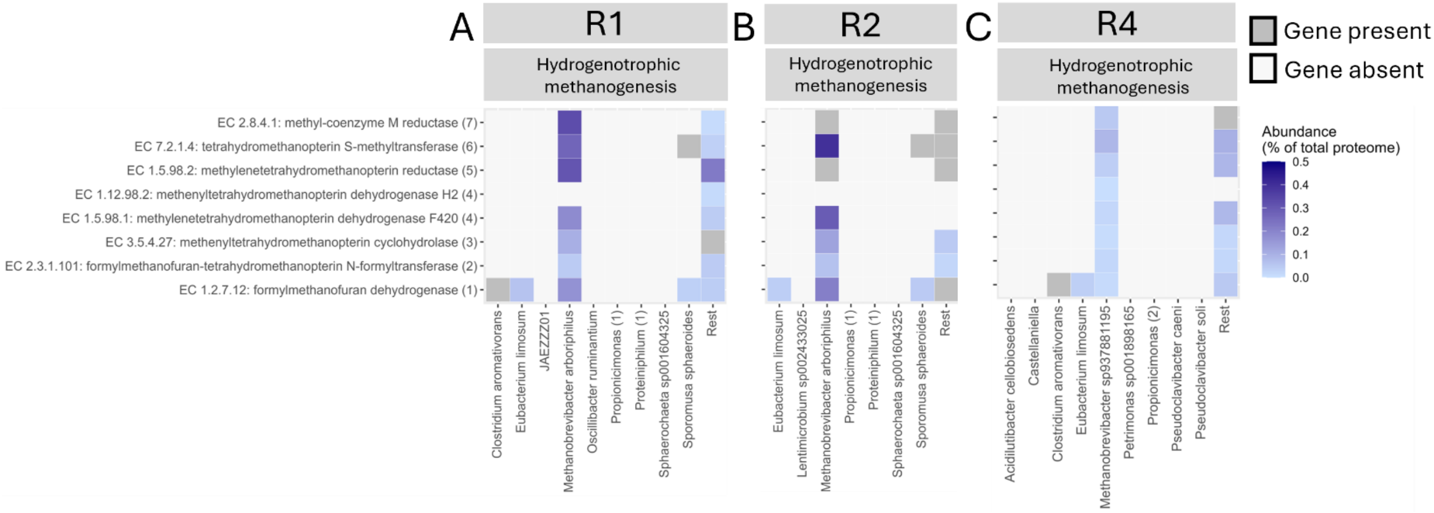


*Figure S14. Heatmaps showing the gene presence and protein abundance of the individual steps of hydrogenotrophic methanogenesis per MAG in reactor 1 (A), reactor 2 (B) and reactor 4 (C). Gene presence is shown in dark grey and protein abundance (%) in light to dark blue gradient.*

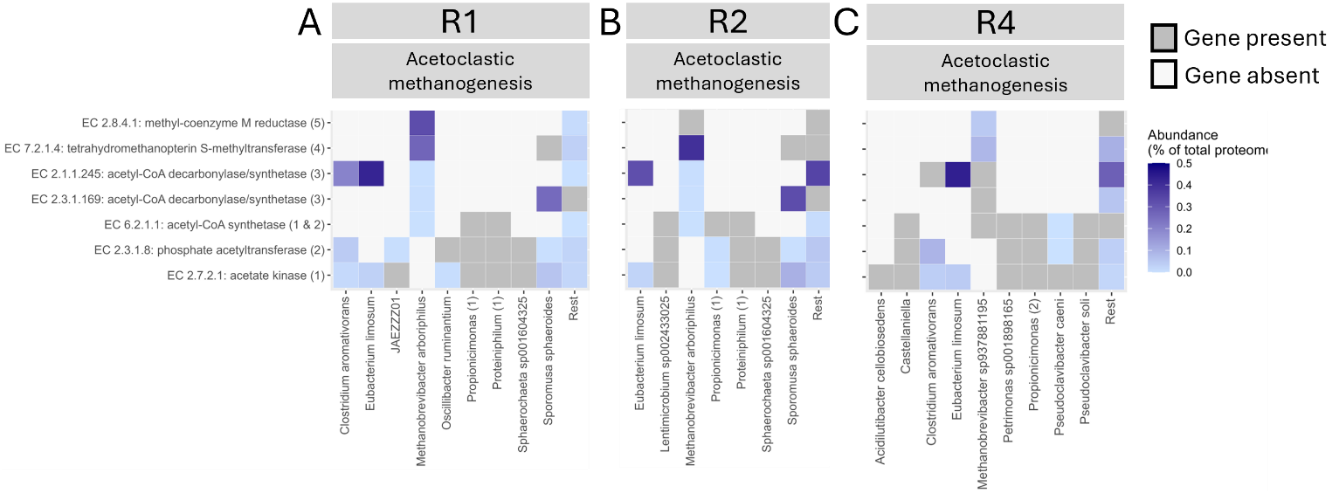


*Figure S15. Heatmaps showing the gene presence and protein abundance of the individual steps of acetoclastic methanogenesis per MAG in reactor 1 (A), reactor 2 (B) and reactor 4 (C). Gene presence is shown in dark grey and protein abundance (%) in light to dark blue gradient.*

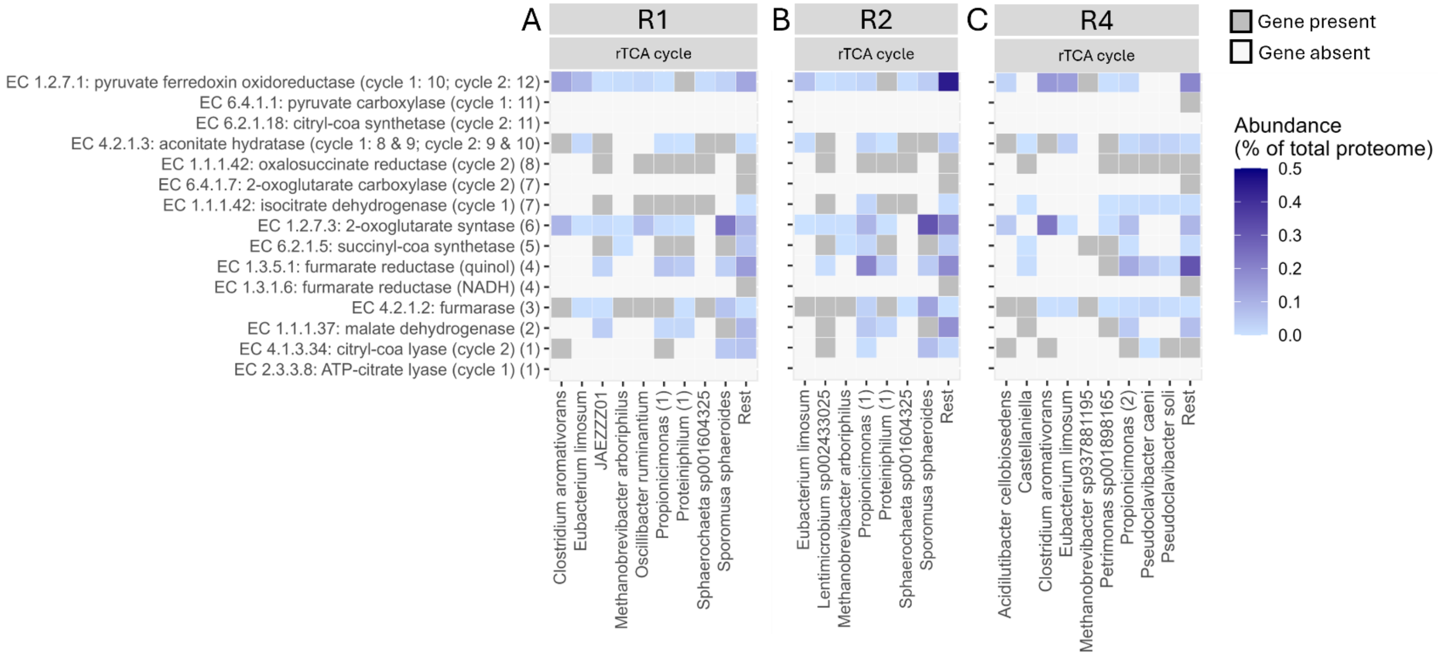


*Figure S16. Heatmaps showing the gene presence and protein abundance of the individual steps of the reverse tri-carboxylic acid cycle (rTCA) cycle per MAG in reactor 1 (A), reactor 2 (B) and reactor 4 (C). Gene presence is shown in dark grey and protein abundance (%) in light to dark blue gradient.*

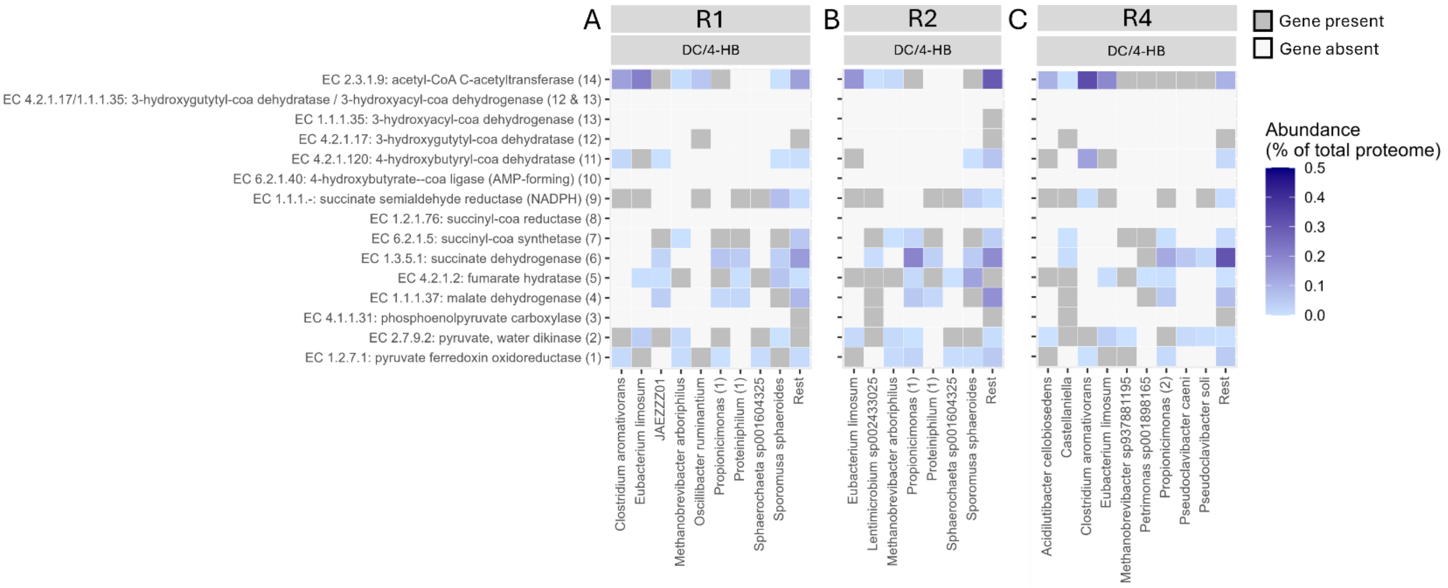


*Figure S17. Heatmaps showing the gene presence and protein abundance of the individual steps of the dicarboxylate-4-hdyroxybutyrate cycle (DC/4-HB) per MAG in reactor 1 (A), reactor 2 (B) and reactor 4 (C). Gene presence is shown in dark grey and protein abundance (%) in light to dark blue gradient.*

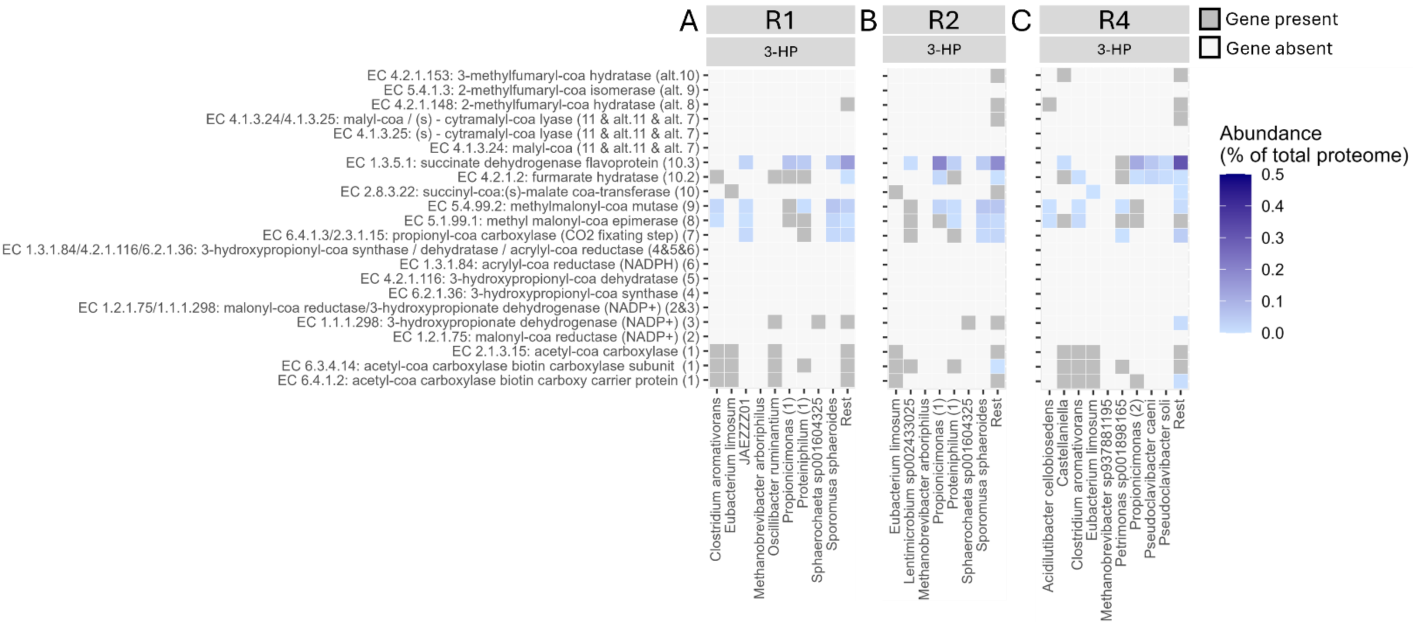


*Figure S18. Heatmaps showing the gene presence and protein abundance of the individual steps of the 3-hydroxypropionate bicycle (3-HP) per MAG in reactor 1 (A), reactor 2 (B) and reactor 4 (C). Gene presence is shown in dark grey and protein abundance (%) in light to dark blue gradient.*

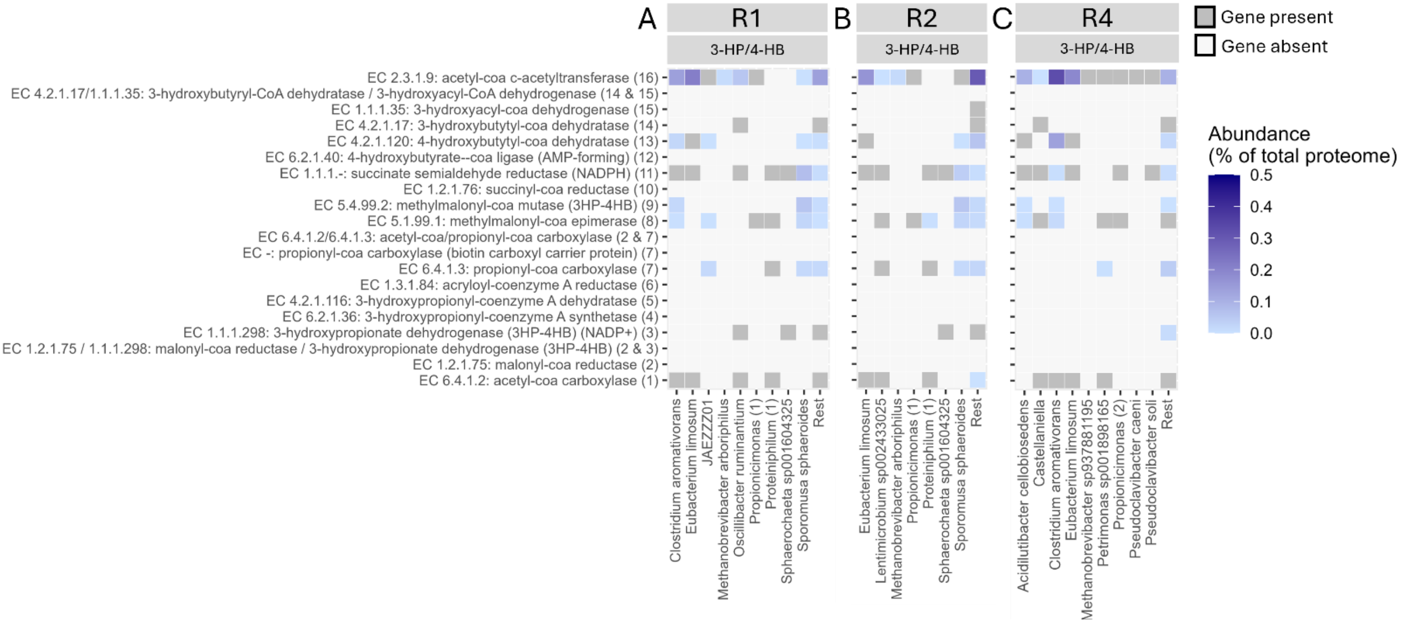


*Figure S19. Heatmaps showing the gene presence and protein abundance of the individual steps of the 3-Hydroxypropionate-4-hydroxybutyrate cycle (3-HP/4-HB) per MAG in reactor 1 (A), reactor 2 (B) and reactor 4 (C). Gene presence is shown in dark grey and protein abundance (%) in light to dark blue gradient.*

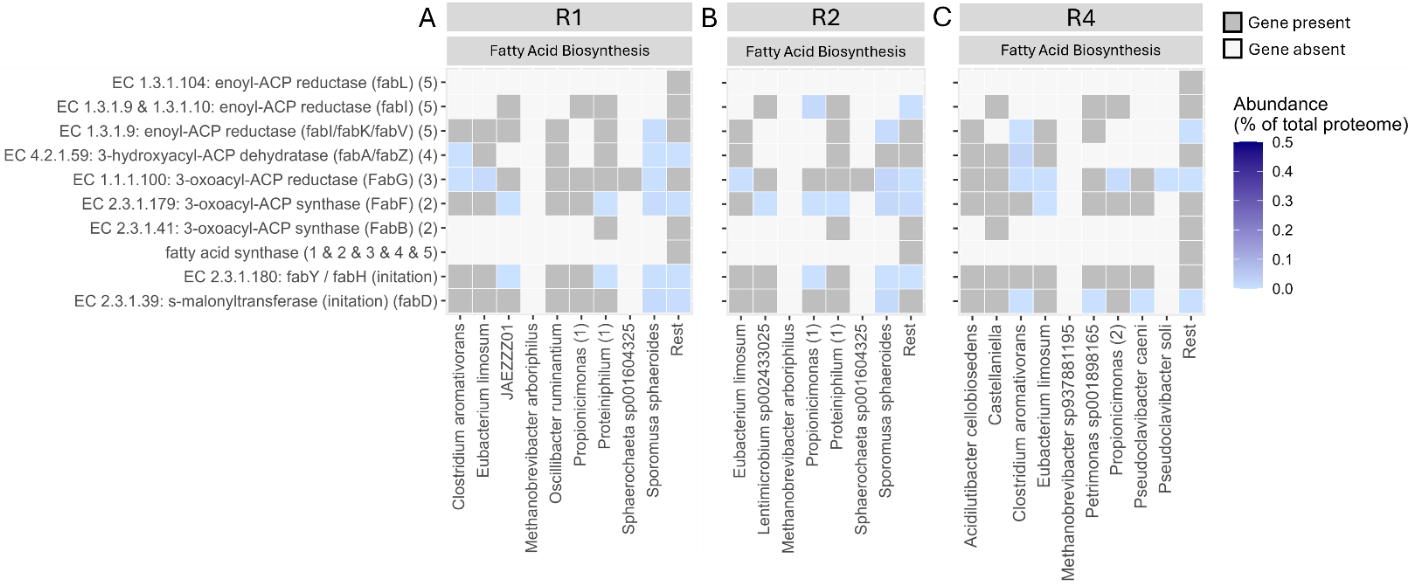


*Figure S20. Heatmaps showing the gene presence and protein abundance of the individual steps of the fatty acid biosynthesis (FAB) per MAG in reactor 1 (A), reactor 2 (B) and reactor 4 (C). Gene presence is shown in dark grey and protein abundance (%) in light to dark blue gradient.*

#### 10. Hypothetical cross-feeding interactions

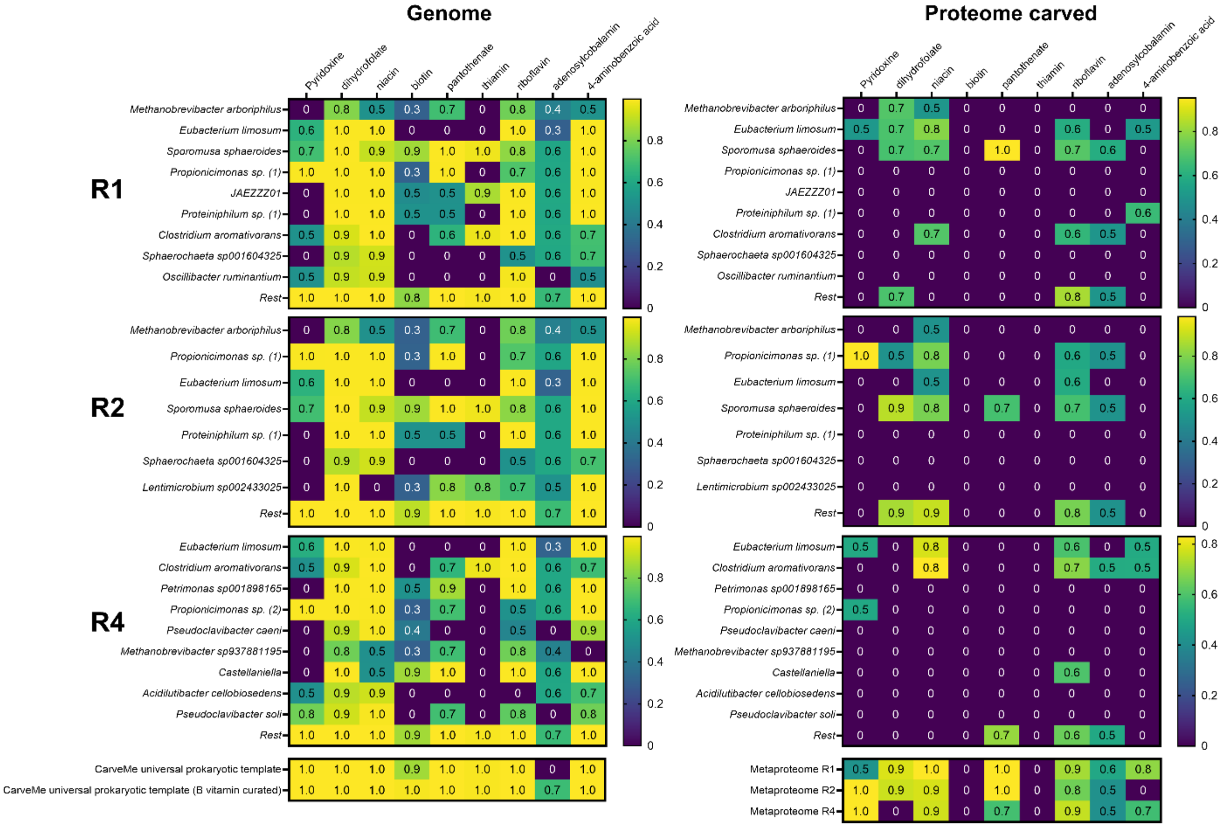


*Figure S21. Producibility metric (PM) values calculated from the metabolic networks constructed from the isolated MAGs in reactor 1, reactor 2 and reactor 4 using the algorithm of Bernstein et al., 2019. PM values were calculated for nine different B vitamins, including pyridoxine, dihydrofolate, niacin, biotin, pantothenate, thiamine, riboflavin, adenosylcobalamin and 4-aminobenzoic acid. The left panel includes the PM values of the metabolic networks that were constructed from the metabolic genes in the genomes. It also includes the PM values calculated for the carveMe universal template that is used for reconstruction and the carveMe universal template curated with the biosynthetic pathways of biotin and adenosylcobalamin. The right panel shows the PM values for the metabolic networks that were carved with the proteome. The proteome carved networks only include those metabolic reactions catalysed by the enzymes that were detected in the proteome. It also includes mixedbag models of the complete metaproteome of each reactor to show the overall producibility capacity based on the proteome.*

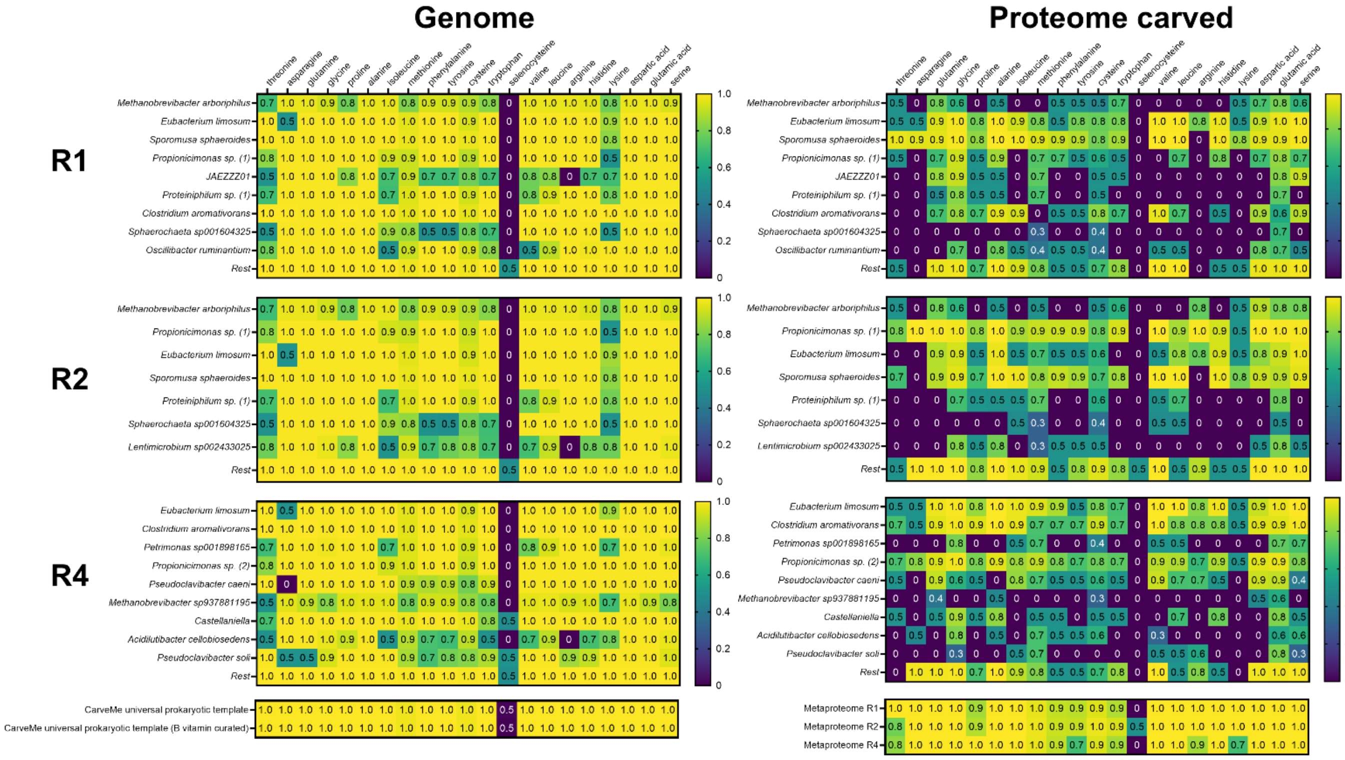


*Figure S22. Producibility metric (PM) values calculated from the metabolic networks constructed from the isolated MAGs in reactor 1, reactor 2 and reactor 4 using the algorithm of Bernstein et al., 2019. PM values were calculated for 21 amino acids, including selenocysteine. The left panel includes the PM values of the metabolic networks that were constructed from the metabolic genes in the genomes. It also includes the PM values calculated for the carveMe universal template that is used for reconstruction and the carveMe universal template curated with the biosynthetic pathways of biotin and adenosylcobalamin. The right panel shows the PM values for the metabolic networks that were carved with the proteome. The proteome carved networks only include those metabolic reactions catalysed by the enzymes that were detected in the proteome. It also includes mixedbag models of the complete metaproteome of each reactor to show the overall producibility capacity based on the proteome.*
